# Human neural stem cells restore spatial memory in a transgenic Alzheimer’s disease mouse model by an immunomodulating mechanism

**DOI:** 10.1101/2023.11.01.565161

**Authors:** Kevin S. Chen, Mohamed H. Noureldein, Lisa M. McGinley, John M. Hayes, Diana M. Rigan, Jacquelin F. Kwentus, Shayna N. Mason, Faye E. Mendelson, Masha G. Savelieffd, Eva L. Feldman

## Abstract

**INTRODUCTION:** Stem cells are a promising therapeutic in Alzheimer’s disease (AD) given the complex pathophysiologic pathways involved. However, the therapeutic mechanisms of stem cells remain unclear. Here, we used spatial transcriptomics to elucidate therapeutic mechanisms of human neural stem cells (hNSCs) in an animal model of AD.

**METHODS:** hNSCs were transplanted into the fimbria fornix of the hippocampus using the 5XFAD mouse model. Spatial memory was assessed by Morris water maze. Amyloid plaque burden was quantified. Spatial transcriptomics was performed and differentially expressed genes (DEGs) identified both globally and within the hippocampus. Subsequent pathway enrichment and ligand-receptor network analysis was performed.

**RESULTS:** hNSC transplantation restored learning curves of 5XFAD mice. However, there were no changes in amyloid plaque burden. Spatial transcriptomics showed 1061 DEGs normalized in hippocampal subregions. Plaque induced genes in microglia, along with populations of stage 1 and stage 2 disease associated microglia (DAM), were normalized upon hNSC transplantation. Pathologic signaling between hippocampus and DAM was also restored.

**DISCUSSION:** hNSCs normalized many dysregulated genes, although this was not mediated by a change in amyloid plaque levels. Rather, hNSCs appear to exert beneficial effects in part by modulating microglia-mediated neuroinflammation and signaling in AD.

## INTRODUCTION

Alzheimer’s disease (AD) is the most common neurodegenerative disorder and frequent cause of dementia (Scheltens et al., 2021). Patients exhibit progressive memory loss, spatial disorientation, and an inability to perform activities of daily living. Pathologically, the disease is characterized by an accumulation of amyloid-beta (Aβ) plaques and neurofibrillary tangles of aggregated tau protein (Savelieff et al., 2013), which are linked to neuronal degeneration, particularly in the hippocampus, entorhinal cortex, and eventually throughout the cerebral cortex (Brettschneider et al., 2015). Nevertheless, it remains unclear whether Aβ and tau are causative factors in AD pathogenesis or are simply markers of other underlying pathologic processes. Indeed, AD is increasingly thought to proceed through a multifactorial pathology, involving neuroinflammation (Heneka et al., 2015; Arranz and De Strooper, 2019), disrupted metabolism (Serrano-Pozo et al., 2021), and vascular dysfunction, in addition to Aβ and tau pathology (Scheltens et al., 2021).

AD still lacks effective disease-modifying treatments, due, in part, to this complex pathophysiology involving multiple disrupted pathways. Targeted pharmaceutical interventions have previously failed to reverse or even arrest disease progression (Shi et al., 2020), which has prompted investigation into therapeutics designed to impact alternative or multiple pathologic processes, such as neuroinflammation. Stem cell-based approaches are especially attractive in this regard because they can enrich the nervous system milieu in a sustained fashion through multiple biological pathways, including immunomodulatory, neuroprotective, metabolic, toxin scavenging, and other mechanisms (Goutman et al., 2019; Boese et al., 2020; Sakowski and Chen, 2022).

We previously showed that a human neural stem cell line (hNSC) improves spatial memory in an AD mouse model (McGinley et al., 2018). These hNSCs express insulin-like growth factor-I (IGF-I), survive in the long-term (McGinley et al., 2022), and exert immunomodulatory (McGinley et al., 2018) and neuroprotective (McGinley et al., 2016) effects in the brain of AD mice following transplantation into the fimbria fornix of the hippocampus. In the current study, we performed spatial transcriptomics of brain tissue following hNSC transplantation into transgenic 5XFAD AD mice. Our goal was to understand the neuroprotective properties and underlying mechanisms of transplanted stem cells. We report that hNSC transplantation improves spatial memory in 5XFAD mice. Spatial transcriptomics reveals hNSCs induce local effects in multiple brain regions of 5XFAD mice by normalizing amyloid plaque-related gene expression, notably in microglial/inflammatory pathways. Our findings confirm that hNSC transplantation exerts beneficial effects in the brain of AD mice, mediated, in part, by modulating inflammatory pathways in the brain.

## METHODS

### Animal model

All animal procedures were approved by the University of Michigan Institutional Animal Care and Use Committee (Protocol #PRO00010247). The 5XFAD transgenic mouse model was used (catalog # 034848, The Jackson Laboratory, Bar Harbor, ME), expressing five mutations associated with familial AD: the Swedish (K670N/M671L), Florida (I716V), and London (V717I) mutations in the human amyloid-beta precursor protein (APP), as well as human presenilin-1 (PS1) harboring M146L and L286V mutations.

### hNSC transplantation and immunosuppression regimen

An IGF-1 producing hNSC cell line (Palisade Bio, Carlsbad, CA) was cultured as described previously (McGinley et al., 2016; McGinley et al., 2018). On the day of the transplantation procedure, hNSCs were trypsinized, centrifuged, and resuspended in hibernation media (Seneca Biopharma) and stereotactically transplanted as previously described (McGinley et al., 2018). Briefly, animals were anesthetized with 1-2% isoflurane inhaled on a 100% oxygen carrier, positioned within a mouse cranial stereotactic frame in the standard manner, and a powered burr was used to create a skull opening large enough to permit introducing a 34-gauge needle. Six infusions in total of 1 μL each were targeted to three sites bilaterally in the fimbria fornix of the hippocampus at the following coordinates relative to the bregma (millimeters posterior/lateral/ventral): 0.82/0.75/2.5, 1.46/2.3/2.9, and 1.94/2.8/2.9. Each infusion was administered over 60 seconds, followed by a 60 second equilibration period before withdrawing the needle and closing the scalp.

For cohorts undergoing behavioral testing and histological/ELISA Aβ analysis, mice were randomly assigned to four treatment groups at 26 weeks of age: (i) untreated non-transgenic wild-type mice (WT) littermates (n=10), (ii) untreated 5XFAD (n=10), (ii) vehicle injected 5XFAD (n=9), and (iii) hNSC transplanted 5XFAD (n=10). Vehicle animals underwent injections of hibernation media. For hNSC intracranial transplantation, 3 bilateral injections of hNSCs at 30,000 cells/µL, 1 µL per injection for a total 180,000 cells were performed per animal. Procedural animals received mycophenolate mofetil (30 mg/kg subcutaneous) daily for 7 days after the procedure, and tacrolimus (3.0 mg/kg subcutaneous) was administered daily starting at 24 weeks of age until study end.

Brain sections utilized for spatial transcriptomics were obtained from WT mice, untreated 5XFAD mice, 5XFAD mice receiving immunosuppression alone, and 5XFAD mice undergoing intracranial hNSC transplantation. Here, immunosuppression consisted of monoclonal antibodies against mouse CD4 (clone GK1.5, rat IgG2b,κ; catalog # BE0003-1, Bio X Cell, Lebanon, NH) and CD40L (clone MR-1, Armenian hamster IgG; catalog # BE0017-1, Bio X Cell). Injections (20 mg/kg intraperitoneal for each antibody) began starting 1 day prior to the surgical procedure and occurred daily for 4 days, then weekly thereafter for the duration of the experiment. hNSC were transduced to express GFP and firefly luciferase (McGinley et al., 2022), and transplants were performed in 5XFAD mice at 6 weeks of age (3 bilateral injections of hNSCs at 50,000 cells/µL, 2 µL per injection, for a total of 600,000 cells per animal).

### Morris water maze

The Morris water maze is a well-established method of assessing spatial memory in rodent models (Bromley-Brits et al., 2011). Briefly, at 34 to 35 weeks of age (8 weeks after hNSC or vehicle injection for procedural groups), animals were placed in a pool of opaque water with visual cues placed around the perimeter of the pool. A submerged hidden platform was placed in one quadrant, which provides an escape from water. The mice can deduce the location of the hidden platform from the spatial relationship to the surrounding visual cues. With intact short-term memory, the latency for animals to swim towards and find the hidden platform decreases over repeated trials as animals learn the spatial relationship between the platform and visual cues (4 trials per day for 9 days). A positive control with a visible platform was performed on the 10^th^ day. Subsequently, to test long-term reference memory, animals were reintroduced into the maze 7 days after the last trial, except the platform was removed as a probe trial. Time spent probing each quadrant of the pool was measured. Mice with intact spatial learning and long-term memory will spend a disproportionate amount of time searching for the platform in the quadrant where it was previously placed. On the other hand, mice with impaired spatial memory spend only 25% of the time in the correct quadrant by chance.

### Tissue preparation and immunohistochemistry

After completing behavioral testing, animals are euthanized at 35 weeks of age by phenobarbital overdose and transcardially perfused with phosphate buffered saline (PBS) solution. Brains were dissected from the calvarium and bisected along the midline sagittal plane. One brain hemisphere was fixed in 4% paraformaldehyde for 24 h, cryoprotected in escalating sucrose gradients, embedded, and cryosectioned onto histologic slides (10 μm thickness, coronal plane sections). Hematoxylin/eosin staining was performed on every 10^th^ slide to identify hippocampal structures and the approximate location of hNSC injections.

For immunohistochemistry slides were warmed and washed with PBS, then permeabilized/blocked with 0.3% Triton X-100 and 5% bovine serum albumin (BSA). Slides were then incubated in primary rabbit anti-Aβ antibody (1:1500; catalog # 9888, Cell Signaling Technologies, Danvers, MA) diluted in 0.1% Triton X-100 and 5% BSA overnight at 4 °C. Sections were washed with PBS and incubated with fluorescent anti-rabbit secondary antibody at 1:1000 in 0.1% Triton X-100 and 5% BSA for 1 h. After washing in PBS again, slides were incubated with Hoechst nuclear stain (1 mg/mL) for 10 min, washed, and mounted with glass coverslips and Prolong Gold Anti-Fade mountant (Thermo Fisher Scientific, Waltham, MA).

Fluorescent images were captured at 20X magnification (Nikon Microphot-FXA, Nikon, Chiyoda, Japan; CellSens Dimension software, Olympus, Shinjuku, Japan). Target regions included images of dentate gyrus, cornu ammonis fields, and fimbria fornix. Five images per section were captured from two sections per animal for a total of 10 images per animal. Images were converted to 8-bit and the image thresholds were standardized in Fiji/ImageJ (Schindelin et al., 2015). Images were then processed using the Analyze Particles function to quantify the percentage area of immunoreactivity in each section (plaque-to-plaque free area).

### Multiplex ELISA

The remaining brain hemisphere from each animal was flash frozen in liquid nitrogen and stored at -80°C. Production of AD-related and immune-related factors was assessed by multiplex ELISA (Eve Technologies, Calgary, Canada). Brain tissue was harvested after Morris water maze testing, thawed on ice and lysed in RIPA buffer (catalog # 89900, Thermo Fisher Scientific) containing a protease inhibitor cocktail (catalog # 11697498001, Roche Diagnostics, Basel, Switzerland). Lysates were collected, briefly sonicated, brought to a volume of 1 mL and centrifuged at 13,000 rpm for 20 min at 4 °C. Supernatants were collected, diluted ten-fold in RIPA buffer and protein concentration was measured and normalized to 400 µg/mL. Samples were flash frozen in liquid nitrogen and shipped on dry ice to Eve Technologies for analysis. Discovery Assay protein arrays and Custom-Plex Assays were used to quantify the following analytes: Human Aβ and Tau 2-Plex Assay (custom panel: Aβ42, Aβ40), Human Supplemental Biomarker 10-Plex Discovery Assay [catalog # HDHSB10: neural cell adhesion molecule (NCAM), soluble vascular cell adhesion molecule 1 (sVCAM-1), soluble intercellular adhesion molecule 1 (sICAM-1), platelet-derived growth factor (PDGF-AA, PDGF-AB), cathepsin D], mouse matrix metalloproteinase (MMP) 5-Plex Discovery Assay (catalog # MDMMP-S,P: MMP-2, MMP-3, MMP-8, MMP-9, MMP-12), mouse transforming growth factor beta (TGF-β) 3-Plex Discovery Assay (catalog # TGFβ1-3: TGF-β1, TGF-β2, TGF-β3), and Mouse Focused 10-Plex Discovery Assay [catalog # MDF10: interleukins (IL-1β, IL-2, IL-6, IL-10, IL-12), granulocyte-macrophage colony-stimulating factor (GM-CSF), monocyte chemoattractant protein 1 (MCP-1), and tumor necrosis factor alpha (TNF-α)].

### Statistics

Behavioral data was analyzed by two-way analysis of variance (ANOVA), and probe trial was analyzed by one-way ANOVA using Prism (GraphPad, La Jolla, CA). For immunohistochemistry and ELISA data, statistical differences were determined by one-way ANOVA. A p<0.05 was considered statistically significant and all values are presented as mean ± SEM (standard error of the mean). Spatial transcriptomics data were analyzed using R version 3.5.1 and RStudio 1.3.1093.

### Spatial transcriptomics

In a separate experiment for spatial transcriptomics of the brain, we compared hNSC injected 5XFAD animals to 5XFAD mice receiving immunosuppression alone, untreated 5XFAD mice, and control WT mice. At 34 weeks post-transplantation, all animals were euthanized and perfused with PBS on the same day. Brains were dissected, hemisected along the sagittal midline, placed in OCT media, frozen in isopentane cooled with liquid nitrogen, and stored at -80 °C. Test coronal sections, 10 μm thick, were placed on Visium 10X tissue optimization slides (10X Genomics, Pleasanton, CA) to optimize permeabilization conditions.

Next, all experimental hemibrains were removed from -80 °C storage and thawed on the same day, such that they were only subjected to a single freeze/thaw cycle. All hemibrains were sectioned to 10 μm thick coronal sections, which were placed on Visium 10X slides and submitted simultaneously to the University of Michigan Advanced Genomics Core for analysis. There were 2 sections from WT brain from 1 animal, 4 sections from 5XFAD brain from 2 animals, 2 sections from immunosuppression alone brain from 1 animal, and 6 sections from hNSC transplanted brain from 3 animals. All Visium slides were processed simultaneously over two days according to the manufacturer’s instructions. Briefly, slides were heated at 37 °C for 1 min on a T100 thermal cycler (BioRad, Hercules, CA), transferred to precooled methanol (Sigma Aldrich, St. Louis, MO) for 30 min, and stained by hematoxylin and eosin (H&E) using a standard protocol. Brightfield images of the H&E-stained sections were captured using an AxioObserver (Zeiss, Oberkochen, Germany). Tissue sections were permeabilized for 12 min, as optimized on test brain sections, using the Visium Spatial Tissue Optimization Slide & Reagent kit. Permeabilization releases mRNA from cells, which bind to oligonucleotides on the capture areas followed by reverse transcription, second-strand synthesis, denaturation, and cDNA amplification, as specified in the T100 thermal cycler protocol using the spatial slide adaptor. Amplified cDNA clean-up was performed using SPRIselect (Beckman Coulter, Brea, CA) and the library was prepared according to the manufacturer’s protocol. The generated cDNA libraries were subjected to 151 bp paired-end sequencing, according to the manufacturer’s protocol (NovaSeq 6000, Illumina, San Diego, Ca).

### Data and Image Analysis for Spatial Transcriptomics

For the spatial transcriptomics data, the NovaSeq 6000 sequencer generated “bcl” files, which were converted to de-multiplexed “fastq” files using Bcl2fastq2 Conversion Software (Illumina, San Diego, CA). Space Ranger 1.3.1 (10X Genomics) aligned and filtered reads and counted barcodes and unique molecular identifiers, using the “Fastq” files and H&E image tiff files as inputs. Reads were aligned against the Genome Reference Consortium Mouse Build 38 patch release 6 (GRCm38.p6). Space Ranger correlates the spatial coordinates to gene expression data by reallocating the barcodes in each read to the respective 55 µm feature (*i.e*., single spots) relative to the slide’s fiducial frame. Spatial transcriptomics data were analyzed using R version 4.1.0 and RStudio 1.4.1717 on the Greatlakes cluster, available at the University of Michigan (https://arc.umich.edu/greatlakes/). Preprocessed sequencing data were analyzed with the Seurat R package (version 4.0.5) (Satija et al., 2015) following the guided tutorial pipeline (available at https://satijalab.org/seurat/articles/spatial_vignette.html, accessed on 5/27/2022). Individual datasets were normalized using the “SCTransform” function in the Seurat package by regularized negative binomial regression (Hafemeister and Satija, 2019). SCTransformation was specifically performed to account for heterogeneity in single-cell RNA-seq due to batch effects and experimental variation without losing biological signal.(Choudhary and Satija, 2022) Individual section datasets were integrated using “FindIntegrationAnchors” on the top 3,000 most variable genes, and the “IntegrateData” function from the Seurat package compared the different conditions (Butler et al., 2018).

### Detecting hNSCs

To detect hNSCs in brain sections spatial transcriptomics data and identify their differentially expressed genes, the raw sequencing “fastq” files were realigned to the human genome. As hNSCs were tagged with emerald GFP (EmGFP; https://www.snapgene.com/resources/plasmid-files/?set=fluorescent_protein_genes_and_plasmids&plasmid=Emerald_GFP), the EmGFP sequence was appended to the “Homo_sapiens.GRCh38.ensembl” human genome using the “mkgtf” function in “Spaceranger” to create a Spaceranger-compatible reference genome file. Subsequently, the raw “fastq” files were aligned against this custom reference genome. hNSCs were identified as cells expressing EmGFP and a count matrix of expressed genes was constructed for these cells. A heatmap of the most expressed 100 genes in hNSCs was generated using the pheatmap R package (v 1.0.12) (Kolde, Raivo, “Package ‘pheatmap’.” R package 1, no. 7 (2015): 790).

EmGFP sequence: ATGGTGAGCAAGGGCGAGGAGCTGTTCACCGGGGTGGTGCCCATCCTGGTCGAGCTGGACGGCGACGTAAACGGCCACAAGTTCAGCGTGTCCGGCGAGGGCGAGGGCGATGCCACCTACGGCAAGCTGACCCTGAAGTTCATCTGCACCACCGGCAAGCTGCCCGTGCCCTGGCCCACCCTCGTGACCACCTTGACCTACGGCGTGCAGTGCTTCGCCCGCTACCCCGACCACATGAAGCAGCACGACTTCTTCAAGTCCGCCATGCCCGAAGGCTACGTCCAGGAGCGCACCATCTTCTTCAAGGACGACGGCAACTACAAGACCCGCGCCGAGGTGAAGTTCGAGGGCGACACCCTGGTGAACCGCATCGAGCTGAAGGGCATCGACTTCAAGGAGGACGGCAACATCCTGGGGCACAAGCTGGAGTACAACTACAACAGCCACAAGGTCTATATCACCGCCGACAAGCAGAAGAACGGCATCAAGGTGAACTTCAAGACCCGCCACAACATCGAGGACGGCAGCGTGCAGCTCGCCGACCACTACCAGCAGAACACCCCCATCGGCGACGGCCCCGTGCTGCTGCCCGACAACCACTACCTGAGCACCCAGTCCGCCCTGAGCAAAGACCCCAACGAGAAGCGCGATCACATGGTCCTGCTGGAGTTCGTGACCGCCGCCGGGATCACTCTCGGCATGGACGAGCTGTACAAGTAA

Few GFP-positive hNSC-containing spots were detected, suggesting aligned transcript reads to the mouse genome were likely not contaminated by transcripts from human cells to any significant extent. To confirm this, hNSC sections were also aligned against a mixed mouse/human genome, which found that only 0.9% to 3.9% of sequencing reads from hNSC sections aligned to the human genome. These reads were distributed mainly to GFP-positive spots. A smaller fraction of reads was present in other spots, which mainly contained mouse-aligned reads. Aligning reads to the mixed mouse/human genome did not significantly impact results compared to aligning to the mouse genome.

### Dimension reduction, clustering, and visualization

Principal component analysis was conducted using the top 3,000 most variable genes. Clusters were detected using the first 30 principal components. The number of principal components was selected based on a percent change in variation in consecutive principal components less than 0.1%. Next, Uniform Manifold Approximation and Projection (UMAP) of the principal components was used to visualize cells. Graph-based clustering was performed on the principal component analysis-reduced data.

### Cell type annotation and differential expression analysis

To assign each cluster a cell type identity, cluster gene markers were identified using the “FindAllMarkers” function in Seurat. Clusters were annotated based on gene expression patterns and the H&E images using the Allen Brain Atlas (Lein et al., 2007). After distinct brain regions were annotated, different glia were identified based on the expression of canonical markers as follows, oligodendrocytes (*Mag*, *Mog*, *Olig2*), astrocytes (*Gfap*, *Slc1a3*), microglia (*Itgam*, *Ptprc*), stage 1 disease-associated microglia (DAM) (*Trem2*, *Tyrobp*), and stage 2 DAM (*Cst7*, *Itgax*, *Spp1*) (Garden and Campbell, 2016; Deczkowska et al., 2018). Wilcoxon test compared proportions of different cell types across different conditions. Non-parametric Wilcoxon rank sum test compared gene expression across different conditions and identified differentially expressed genes (Alves and Higdon, 2013). Differentially expressed genes were identified across conditions in whole brain sections as well as hippocampus. Genes were considered differentially expressed if adjusted p-value<0.05. Pathway enrichment was performed using the Kyoto Encyclopedia of Genes and Genomes (Kanehisa and Goto, 2000) and Gene Ontology (Harris et al., 2004) databases by applying the richR package (v 0.0.19, https://github.com/hurlab/richR, accessed on 5/27/2022). Pathways were considered significant if adjusted p-value<0.05.

All hemibrain sections were processed simultaneously on Visium slides for spatial transcriptomics to minimize batch effects. Additionally, sections from each experimental group were distributed across slides. However, to analyze any possible batch effects, a linear mixed effects model (LMM) was performed. LMM adjusts for different slides by treating sections placed on the same slide as dependent observations. The LMM model was constructed with a random intercept (i.e., the slide) and without a random slope using the model formula form ∼condition+(1|slide). The LMM was applied using the lmerSeq R package on the variance-stabilizing transformed pseudobulk aggregated counts.(Vestal et al., 2022) The majority of DEGs from LMM (**Table S14** for whole brain; **Table S15** for hippocampus) overlapped with DEGs by Wilcoxon, suggesting the absence of batch effects and the robustness of the SCTransformation method.

### Cell-to-cell communication

CellChat examined communications across cells (Jin et al., 2021). CellChat leverages network analysis and pattern recognition to predict signaling outputs from cells and signaling inputs to cells. CellChat also analyzes how input and output signals across cells coordinate. First, CellChat identifies significant ligand-receptor pair signaling pathways across all cell clusters. Second, the software predicts incoming signals to and outgoing signals from specific cell clusters. CellChat also predicts global communication patterns using pattern recognition approaches. Similarity measures and manifold learning from topological perspectives organizes signaling pathways. Lastly, CellChat calculates the communication probability of a signaling pathway by summarizing the probabilities of its associated ligand-receptor pairs.

### PROGENy analysis

In addition to richR for pathway enrichment, PROGENy (Pathway RespOnsive GENes) was also used to estimate the activity of relevant signaling pathways based on consensus gene signatures (Schubert et al., 2018). PROGENy leverages publicly available signaling perturbation experiments to yield a common core of 14 pathways comprising genes responsive to the perturbations for human and mouse (Holland et al., 2020). PROGENy has the advantage of overcoming the limitations of conventional pathway enrichment methods, such as overlooking the effects of post-translational modifications on pathway activation. PROGENy scores were computed to infer pathway activities from the spatial transcriptomics dataset using PROGENy R package (v 1.17.3) (https://saezlab.github.io/progeny/index.html, accessed on 5/31/2022).

## RESULTS

### hNSC transplantation improves spatial memory in 5XFAD mice after 8 weeks

In the current study, we used the 5XFAD AD model. In this aggressive model, 5XFAD mice develop brain amyloid plaques, along with neurodegeneration, microgliosis, and cognitive impairment. We transplanted IGF-I expressing hNSCs into the fimbria fornix of the hippocampus of 5XFAD mice aged 26 weeks (**Figure 1A**), employing an immunosuppressive regimen to prevent transplant rejection. Previously, we found that IGF-I expressing hNSCs were more effective against AD pathology *in vitro* compared to the parental line lacking IGF-I (McGinley et al., 2016). Therefore, we opted to perform our spatial transcriptomics analysis using the more effective IGF-I-expressing hNSC line with a view towards supporting future translational clinical trials implementing this cell line. We also demonstrated that injected hNSCs chiefly reside within the fimbria fornix and corpus callosum, but that a fraction migrates throughout the hippocampus, cortex, and other areas of the brain (McGinley et al., 2022).

**Figure 1.**
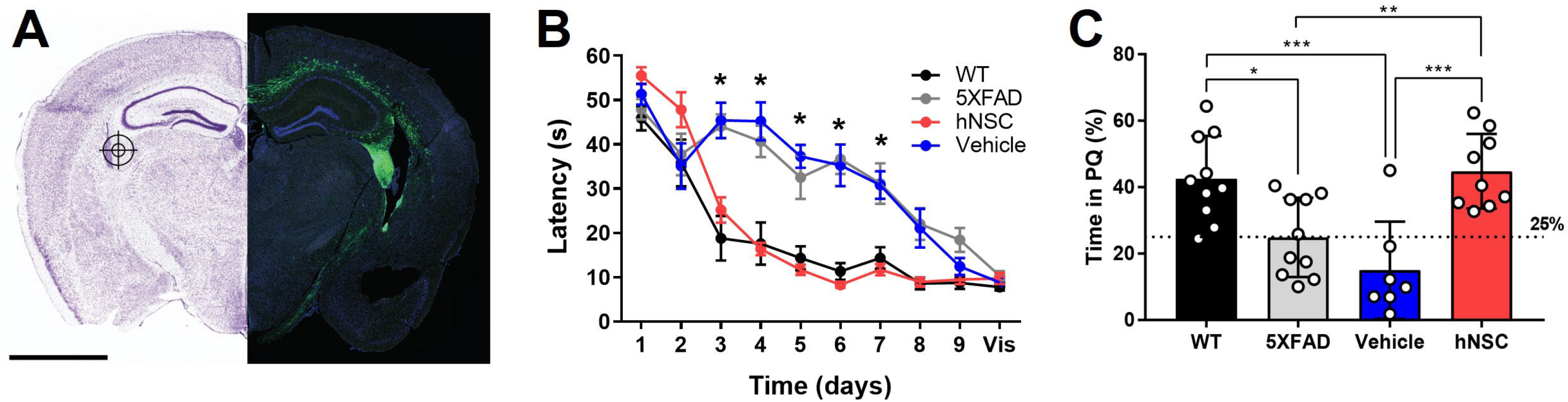
hNSC transplantation improves spatial memory in 5XFAD mice after 8 weeks. (**A**) hNSCs (180,000 cells) or vehicle (hibernation media) were injected at 3 bilateral (6 total) targets into the fimbria fornix (approximate location represented in the figure by a target icon) of 5XFAD mice. Representative fluorescence imaging (right panel; scale bar = 2 mm) shows GFP-labeled hNSCs disperse throughout the cortex following injection. Adapted from (McGinley, Chen et al. 2022). (**B**) Latency to the hidden platform with training over days 1 to 9 and a visible (vis) positive control on day 10 in the Morris Water Maze test for WT (black), untreated 5XFAD (grey), vehicle injected 5XFAD (blue), and hNSC injected 5XFAD (red) mice; n=7-10, *p<0.05, by two-way ANOVA. (**C**) Probe trial on day 7 following completion of hidden platform training in the Morris water maze test for wild-type (WT, black), untreated 5XFAD (grey), vehicle injected 5XFAD (blue), and hNSC injected 5XFAD (red) mice. Horizontal dashed line represents 25% of time spent in the quadrant with the prior hidden platform (PQ), which indicates time spent in the quadrant by chance. n=7-10, *p<0.05, **p<0.01, ***p<0.001, by one-way ANOVA adjusted with Tukey’s multiple comparisons.

Next, we compared hNSC injected animals to vehicle injected 5XFAD mice (immunosuppressive regimen and sham intracranial injection of cell hibernation media only), untreated 5XFAD mice, and control wild-type (WT) mice. We assessed spatial learning and memory by the Morris water maze at 34 weeks of age, 8 weeks following the hNSC/vehicle procedure (**Figure 1B**). As anticipated, untreated 5XFAD animals exhibited impaired spatial learning versus WT controls, with prolonged latencies to the escape platform (p<0.05 on days 3-7; **Figure 1B**). Vehicle injected animals had the same impaired learning curve as 5XFAD animals, demonstrating that neither the immunosuppressive regimen nor the surgical procedure impacts the cognitive phenotype. By contrast, mice receiving hNSCs demonstrated restored learning and memory, and recapitulated the learning curve of WT animals, which did not differ significantly in latency to the platform. All animals were able to reach a visible platform, confirming they did not suffer from any underlying visual, motoric, or motivational deficits.

In the probe trial of long-term memory, performed 7 days after hidden platform training, 5XFAD and vehicle injected animals spent about 25% of their time within the prior platform quadrant (PQ), performing no better than chance (**Figure 1C**). However, hNSC injected animals performed significantly better than vehicle injected animals, spending about 40% of time in the PQ (p<0.0001), like WT animals. Thus, hNSC transplantation into 5XFAD mice restores spatial learning and long-term memory performance to levels commensurate with WT controls.

### hNSC transplantation does not alter brain Aβ levels in 5XFAD mice after 8 weeks

One putative beneficial effect of stem cells in AD is that they lower brain Aβ plaque burden (McGinley et al., 2018; Boese et al., 2020). Thus, we sought to determine whether the beneficial effect of hNSCs on cognitive performance in 5XFAD mice was mediated by direct effects on Aβ levels. In hemispheric brain homogenates, Aβ40 and Aβ42 levels were significantly higher in all transgenic animals versus WT littermates, as assessed by multiplex ELISA (**Figure 2A**). However, hNSC treatment did not change amyloid burden versus vehicle injected animals (Aβ40 hNSC versus vehicle, p=0.91; Aβ42 hNSC versus vehicle, p=0.51). We confirmed this by quantifying Aβ by immunohistochemistry (**Figure 2B**). Substantial plaque pathology was present in all transgenic animals versus WT (p<0.0001; **Figure 2C**). However, plaque load did not differ between hNSC and vehicle injected animals for all imaged regions (p=0.69). This result persisted upon analysis of specific subregions, specifically hippocampal and fimbria fornix structures. Thus, hNSC-induced improvements in spatial learning were not linked to global or brain region-specific alterations in Aβ plaque levels in the aggressive 5XFAD model.

**Figure 2.**
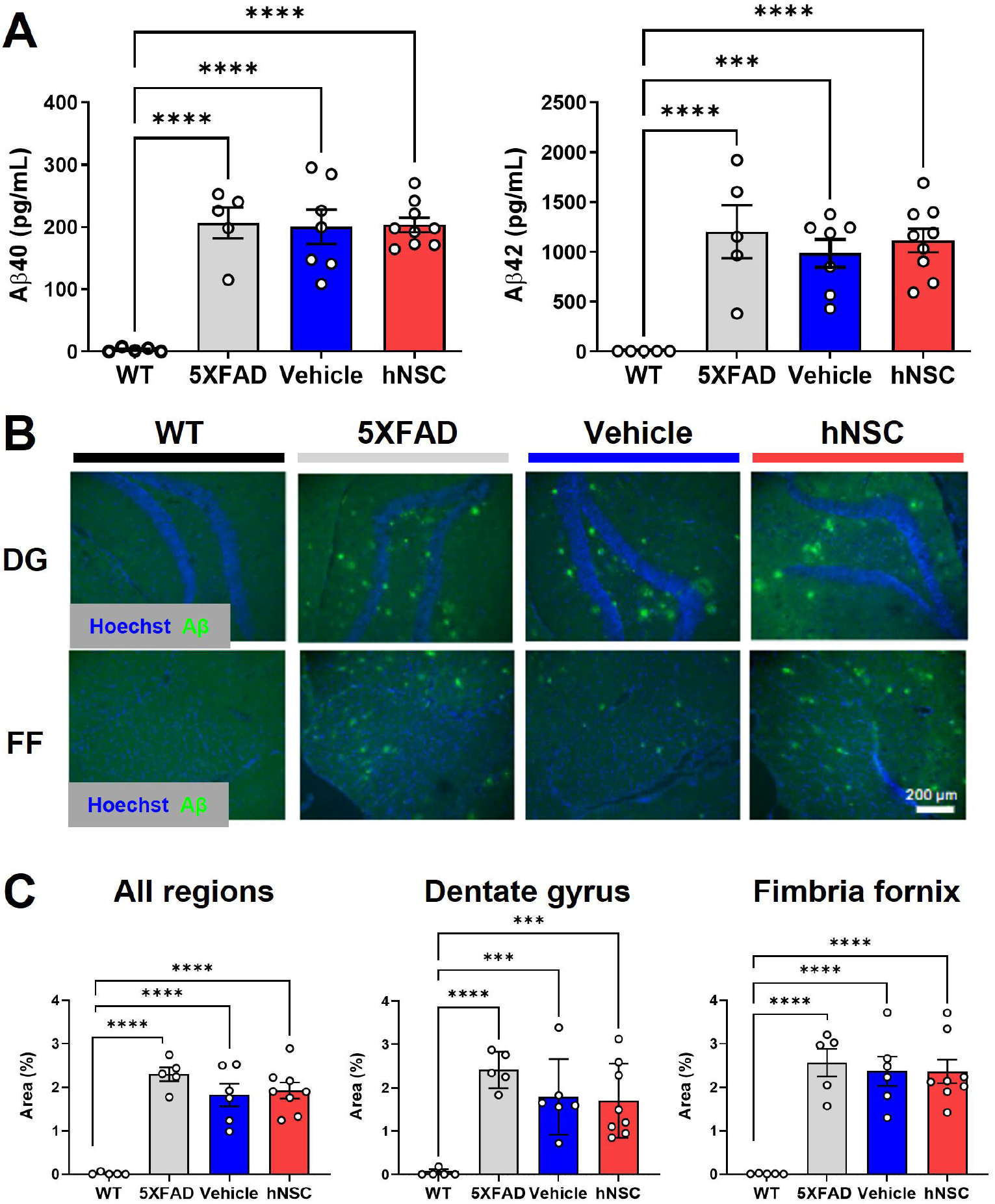
hNSC transplantation does not alter brain Aβ levels in 5XFAD mice after 8 weeks. (**A**) ELISA from brain hemispheres for Aβ40 (left panel) and Aβ42 (right panel) for wild-type (WT, black), untreated 5XFAD (grey), vehicle injected 5XFAD (blue), and hNSC injected 5XFAD (red); n=5-9, ***p<0.001, ****p<0.0001, by one-way ANOVA. (**B**) Representative fluorescence immunohistochemistry image for Aβ (green channel) and Hoechst (blue channel) in the dentate gyrus (DG) and fimbria fornix (FF; scale bar = 200 µm) for WT (black), untreated 5XFAD (grey), vehicle injected 5XFAD (blue), and hNSC injected 5XFAD (red) mice. (**C**) Immunohistochemistry quantification by percent Aβ immunoreactivity area to total area for all brain locations (left panel), dentate gyrus (middle panel), and fimbria fornix (right panel) for WT (black), untreated 5XFAD (grey), vehicle injected 5XFAD (blue), and hNSC injected 5XFAD (red); n=5-8, ***p<0.001, ****p<0.0001, by one-way ANOVA.

### hNSC transplantation does not alter brain cytokine levels in 5XFAD after 8 weeks

A possible mechanism of hNSC-mediated neuroprotection is by interfacing with inflammatory pathways via secreted cytokines (Boese et al., 2020). We previously reported that hNSCs exert an immunomodulatory effect on microglia in APP/PS1 mice (McGinley et al., 2018). We conducted a multiplex ELISA for various cytokines on brain hemisphere homogenates (**Figure S1**). Overall, we noted some variation in brain cytokines, but no changes were specific to hNSC transplants. We also performed a multiplex ELISA assay for various cell adhesion proteins, growth factors, and extracellular matrix remodeling proteins (**Figure S2**). NCAM, sVCAM-1, and cathepsin D (p<0.05 for all) were uniquely elevated in hNSC injected animals compared to vehicle injected animals. Additionally, there were also differential levels in NCAM (p<0.05), sICAM-1 (p<0.01), and PDGF-AB (p<0.05), which were higher in hNSC injected versus non-injected mice. Generally, hNSC transplantation in the 5XFAD mouse may induce some changes in mediators of intercellular adhesion and extracellular matrix remodeling.

### hNSC transplantation is detectable in the brain of 5XFAD mice by spatial transcriptomics after 34 weeks

Because we saw no differences in global amyloid burden 8 weeks post hNSC transplantation, we theorized effects may be more pronounced at a longer timepoint after transplantation (McGinley et al., 2018). Considering future translational studies, effects of stem cells would also need to persist over the long term for significant benefit in humans. To achieve longer-term survival of hNSC transplants, we developed an immunosuppression regimen based on weekly injections of antibodies against CD4 and CD40L (McGinley et al., 2022). We previously demonstrated that hNSCs viably persist in large numbers for over 6 months after transplantation, demonstrating this antibody regimen facilitates long-term graft survival in 5XFAD mice (McGinley et al., 2022).

In this setting, Aβ levels 34-weeks after hNSC transplant, as assessed by quantification of immunohistochemical staining, 5XFAD animals still showed no significant differences across treatment groups, even within subregions of hippocampus (**Figure S3**).

To further elucidate long-term impacts of transplanted hNSCs and given that stem cells likely exert their beneficial effects in a paracrine manner, we sought to apply a spatially-informed analysis across diverse brain areas using spatial transcriptomics. We performed spatial transcriptomics in 5XFAD animals following 34 weeks of hNSC transplantation. Animals received transplants at 6 weeks of age into the fimbria fornix and weekly anti-CD4 and -CD40L injections (McGinley et al., 2022). These animals were compared to WT mice, untreated 5XFAD mice, and 5XFAD mice on immunosuppressants only (IS only).

We first clustered the transcriptomic dataset into cell or region types. Cluster visualization of the spatial transcriptomics dataset by Uniform Manifold Approximation and Projection (UMAP) yielded 25 cell populations within 10 regions, which map to their respective brain anatomy (**Figure S4**). Thus, transcriptional signature-based cell cluster identities accurately reflect their anatomic regions, validating our approach. hNSCs transplanted here express luciferase and GFP; thus, we next leveraged GFP transcripts in the spatial dataset to identify the injected stem cells. As expected, most hNSCs concentrated within the injection area of the fimbria fornix and the neighboring corpus callosum (**Figure 3**); however, there was also evidence the stem cells migrated through white matter tracts to distant structures, including the hippocampus and cerebral cortex. Analysis of hNSC-specific gene expression shows elevated levels of *PLP1*, *EEF1A1*, *FTH1*, *CALM2*, *ATP1B1*, and *SNAP25* transcripts (**Figure S5**). Pathway analysis by both Kyoto Encyclopedia of Genes and Genomes (KEGG) or Gene Ontology (GO) shows a strong enhancement in ribosome, mRNA, and protein synthesis biological processes within hNSCs (**Figure S6**).

**Figure 3.**
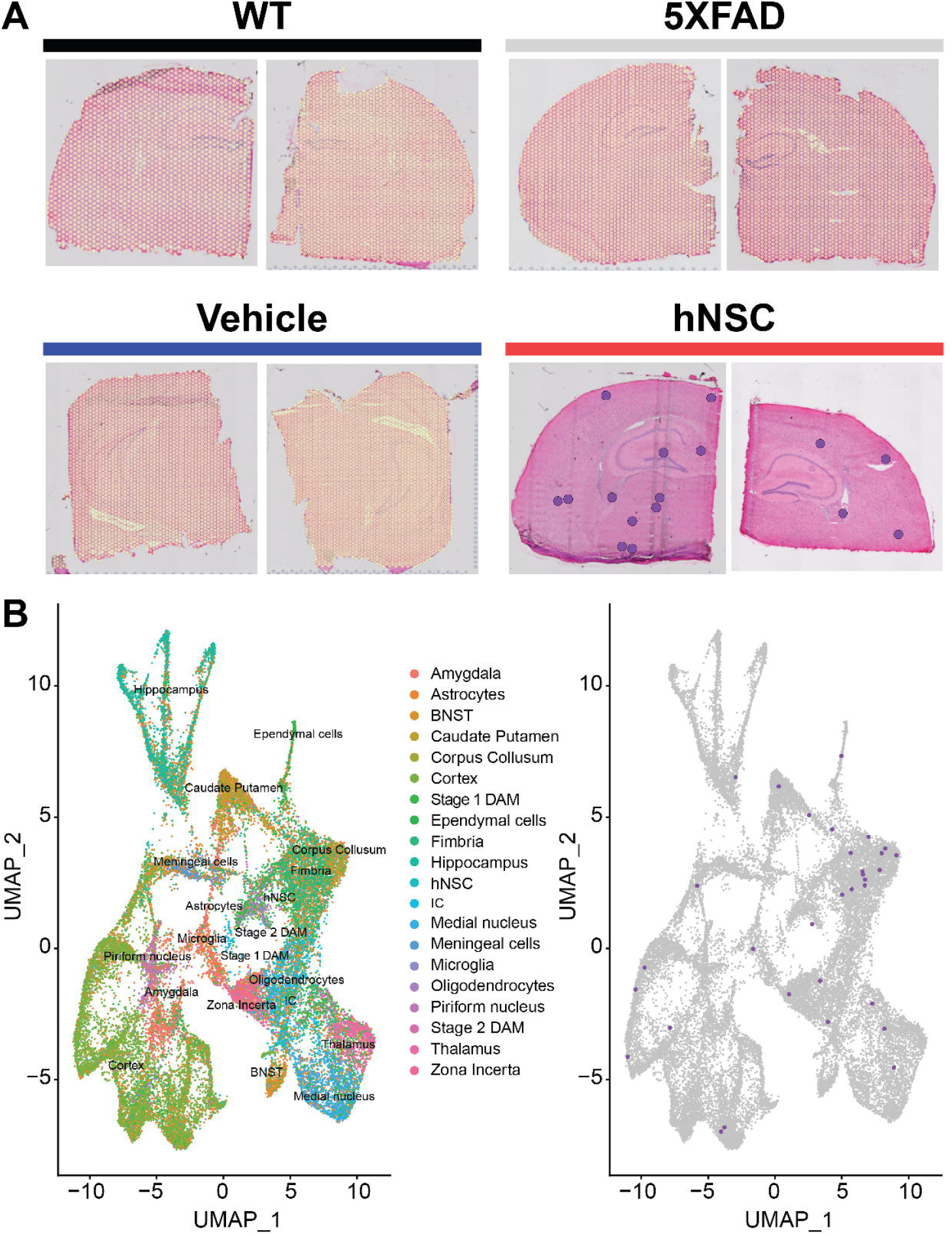
hNSC transplantation is detectable in the brain of 5XFAD mice by spatial transcriptomics after 34 weeks. (**A**) Overlay of detected green fluorescent protein (GFP) transcripts (represented as purple dots) as surrogates of transplanted hNSCs over brain anatomic regions. GFP transcripts were only detected in hNSC injected 5XFAD brain sections (red), but not in untreated 5XFAD (grey), 5XFAD receiving immunosuppression (IS) alone (blue), and wild-type (WT; black) brain sections. Two representative brain sections shown for each condition. (**B**) Left panel: Clustering of the spatial transcriptomics dataset by Uniform Manifold Approximation and Projection (UMAP) yields 25 cell populations (Allen Brain Atlas). Right panel: Overlay of detected GFP transcripts (purple dots) from the spatial transcriptomics dataset against all identified cell clusters. Most GFP transcripts fall within the fimbria fornix and corpus callosum clusters, but are also present within other clusters, e.g., cortex and hippocampus. BNST, bed nucleus of the stria terminalis; DAM, disease associated microglia; hNSC, human neural stem cell; IC, internal capsule.

### hNSC transplantation normalizes differentially expressed genes in brain regions of 5XFAD mice after 34 weeks

After we validated the UMAP-identified cell types against their location in the brain (**Figure S4**), we next examined DEGs, both overall in the brain (**Table S1**) and by cell type or region (**Table S2**). The global brain analysis represents assessment of DEGs based on gene expression across the entire hemisphere section, similar to a bulk analysis. Comparing the untreated 5XFAD model to WT controls, we found there were 2440 upregulated DEGs in 5XFAD versus WT brain, whereas there were only 6 downregulated DEGs in 5XFAD versus WT brain. Next, comparing the treatment effect of hNSCs to the treatment-naïve 5XFAD model, 5052 DEGs were downregulated in hNSC versus 5XFAD (only 2 DEGs were upregulated in hNSC versus 5XFAD). To identify non-hNSC specific perturbations, we found 3037 DEGs were downregulated in IS only versus 5XFAD. Since most DEGs by far were upregulated in 5XFAD mice, we focused on these DEGs, which were subsequently downregulated by hNSC transplantation.

Moreover, we were most interested in 5XFAD DEGs that were specifically normalized by hNSC transplantation, but not altered by IS alone, to gain insight into potential mechanisms of the stem cell-mediated beneficial effects. Thus, we restricted our analysis to hNSC transplantation-specific DEGs by excluding genes that were also affected by IS only. This left 109 DEGs that were significantly and uniquely restored by hNSC treatment, in this global, whole-hemisphere analysis (**Figure S7A**). These 109 DEGs corrected by hNSC transplantation contained several genes known to be related to AD pathogenesis (**Figure S7B**), such as *Trem2* (**Figure S8**), *Tyrobp*, *S100a1*, *Igfbp5*, and complement *C1q*. We next inferred biological significance by performing pathway enrichment analysis of hemispheric DEGs uniquely downregulated by hNSC transplantation versus untreated 5XFAD. Again, pathways related to AD emerged, *e.g*., KEGG terms “lysosome”, “prion disease”, “complement and coagulation cascades”, and several sphingolipid pathways (**Figure S7C**; top panel; **Table S3**), and GO terms “glial cell activation”, “microglial cell activation”, “neuroinflammatory response” (**Figure S7C**; bottom panel; **Table S4**).

### hNSC transplantation normalizes differentially expressed genes in the hippocampus of 5XFAD mice after 34 weeks

Spatial resolution is a significant strength of our dataset. Whereas global brain analysis examined DEGs based on gene expression averaged over the whole brain, region-specific analysis, such as of the hippocampus, reveals spatial heterogeneity in gene expression. Therefore, we next focused on DEG analysis in the hippocampus (**Figure S9**), the limbic structure responsible for short-to long-term memory consolidation, which is adversely impacted in AD. There were 2553 upregulated DEGs in 5XFAD versus WT and 6249 downregulated DEGs in hippocampus from hNSC transplanted animals versus hippocampus from 5XFAD alone. Again, as we were interested in DEGs specifically normalized by hNSCs, we found 1061 were unique to hNSC transplantation after excluding DEGs affected by IS only (**Figure 4A**). There were also 19 downregulated DEGs in 5XFAD versus WT and 1 upregulated DEG in hNSC versus 5XFAD hippocampus, gene “*Bc1*”, which was uniquely impacted by hNSCs.

**Figure 4.**
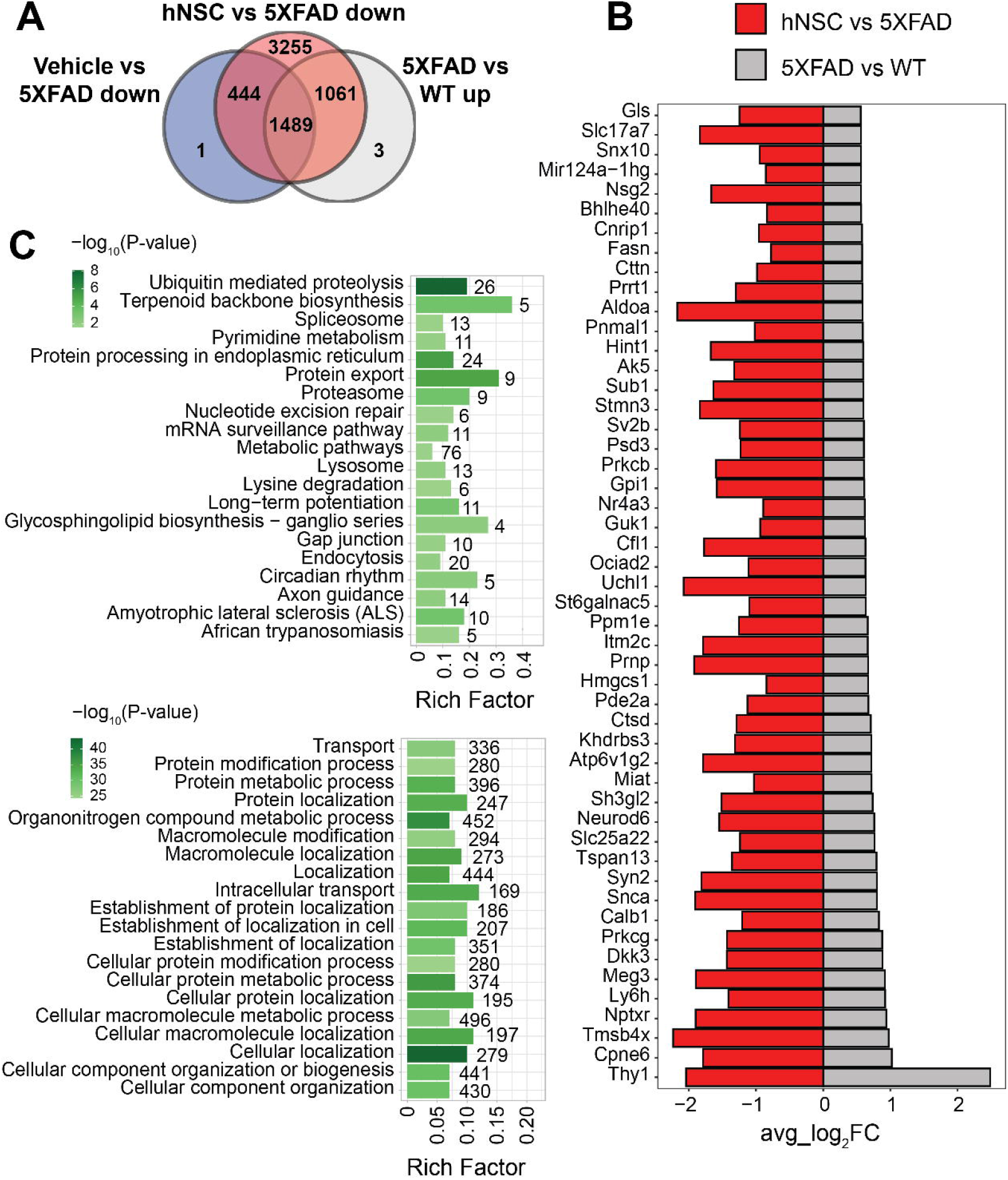
hNSC transplantation normalizes differentially expressed genes (DEGs) in the hippocampus of 5XFAD mice after 34 weeks. (**A**) Venn diagram of upregulated DEGs in 5XFAD versus wild-type (WT; grey) compared to downregulated DEGs in immunosuppressed only (IS only) versus untreated 5XFAD (blue), and hNSC-transplanted versus untreated 5XFAD (red) in hippocampus. Within the subregion of the hippocampus, there were 1061 DEGs upregulated in 5XFAD animals that were subsequently normalized by hNSC treatment (*i.e*., excluding changes seen in IS only animals). (**B**) Bar plot of top 50 DEGs uniquely upregulated in 5XFAD versus WT (grey) and downregulated in hNSC versus 5XFAD (red) in hippocampus. FC, fold-change. Entire DEGs list in **Table S2**. (**C**) Kyoto Encyclopedia of Genes and Genomes (KEGG; top panel) and Gene Ontology (GO; bottom panel) pathway analysis of 1061 DEGs uniquely upregulated in 5XFAD versus WT and downregulated in hNSC versus 5XFAD in hippocampus. Pathway significance represented by color scale from less (light green) to more (dark green) significant. Number over each bar represents the number of DEGs; rich factor represents the proportion of DEGs relative to the total number in the pathway. All these KEGG and GO pathways are activated in 5XFAD mice compared to WT while hNSC (but not IS only) downregulates them all.

The top 50 DEGs upregulated in 5XFAD versus WT hippocampus that were uniquely suppressed by hNSC transplantation contained several DEGs linked to AD, *e.g*., *Neurod6* and *Ctsd* (**Figure 4B**). KEGG analysis of exclusively 5XFAD upregulated and hNSC downregulated DEGs in hippocampus identified pathways involving “ubiquitin mediated proteolysis”, “protein export”, and “proteosome” (**Figure 4C**, top panel; **Table S5**), indicating hNSC transplantation may impact protein degradation dynamics. “Long-term potentiation” was also overrepresented, as was “metabolic pathways”, which contained the largest number of genes at 76. GO analysis likewise found protein and metabolic pathways, such as “protein modification process”, “protein metabolic process”, in addition to highly significant localization pathways, *e.g*., “cellular localization”, “localization” (**Figure 4C**, bottom panel; **Table S6**).

In summary, there were 109 DEGs based on average gene expression across the whole 5XFAD hemibrain that were uniquely reversed by hNSC transplantation (**Figures S6**, **Table S1**). However, when we examined spatial heterogeneity in gene expression, there were 1061 DEGs uniquely rescued by hNSCs when investigation focused within the hippocampus (**Figures 4**, **Table S2**). The presence of more DEGs uniquely reversed by hNSCs in 5XFAD hippocampus versus other brain regions suggests that the stem cells especially ameliorated regions of the brain responsible for memory and learning, highly relevant to AD. This likely resulted from transplantation of the hNSCs into the fimbria fornix of the hippocampus, the white matter outflow tract of the hippocampus; however, our data demonstrate that stem cells also exert a benefit in areas of the brain remote from the transplantation site. Overall, stem cells induce transcriptomic changes across the brain opposing AD pathology in a brain region-specific manner.

### hNSC transplantation normalizes plaque-induced genes and microglial phenotypes in the brain of 5XFAD mice after 34 weeks

In this aggressive 5XFAD AD model, hNSC transplantation for 8 weeks ameliorated cognition, but not bulk changes in Aβ40 and Aβ42 brain levels (**Figure 2**). Thus, in addition to our examination of DEGs in whole brain and hippocampus (**Figures 4**, **S6**), we also analyzed our spatial transcriptomics dataset following 34 weeks of hNSC transplantation to examine local dynamics in amyloid burden. To accomplish this, we leveraged a published panel of plaque-induced genes (PIGs), which specifically associate spatially with Aβ plaques (Chen et al., 2020). This allowed us to assess the impact of stem cells on amyloid burden spatially in fine-grained detail. Of all brain areas or cell types, PIG expression was highest in microglia, across all WT, 5XFAD, IS only, and hNSC conditions (**Figure S10**). Within microglia, PIG expression pattern was relatively low in WT, but increased in the 5XFAD condition (**Figure 5A**). hNSC transplantation rescued the PIGs pattern to one more like control WT brain, whereas IS only did not impact PIGs (**Figure 5A**). Thus, stem cell treatment normalized the expression of genes in microglia that respond to amyloid plaque deposition.

**Figure 5.**
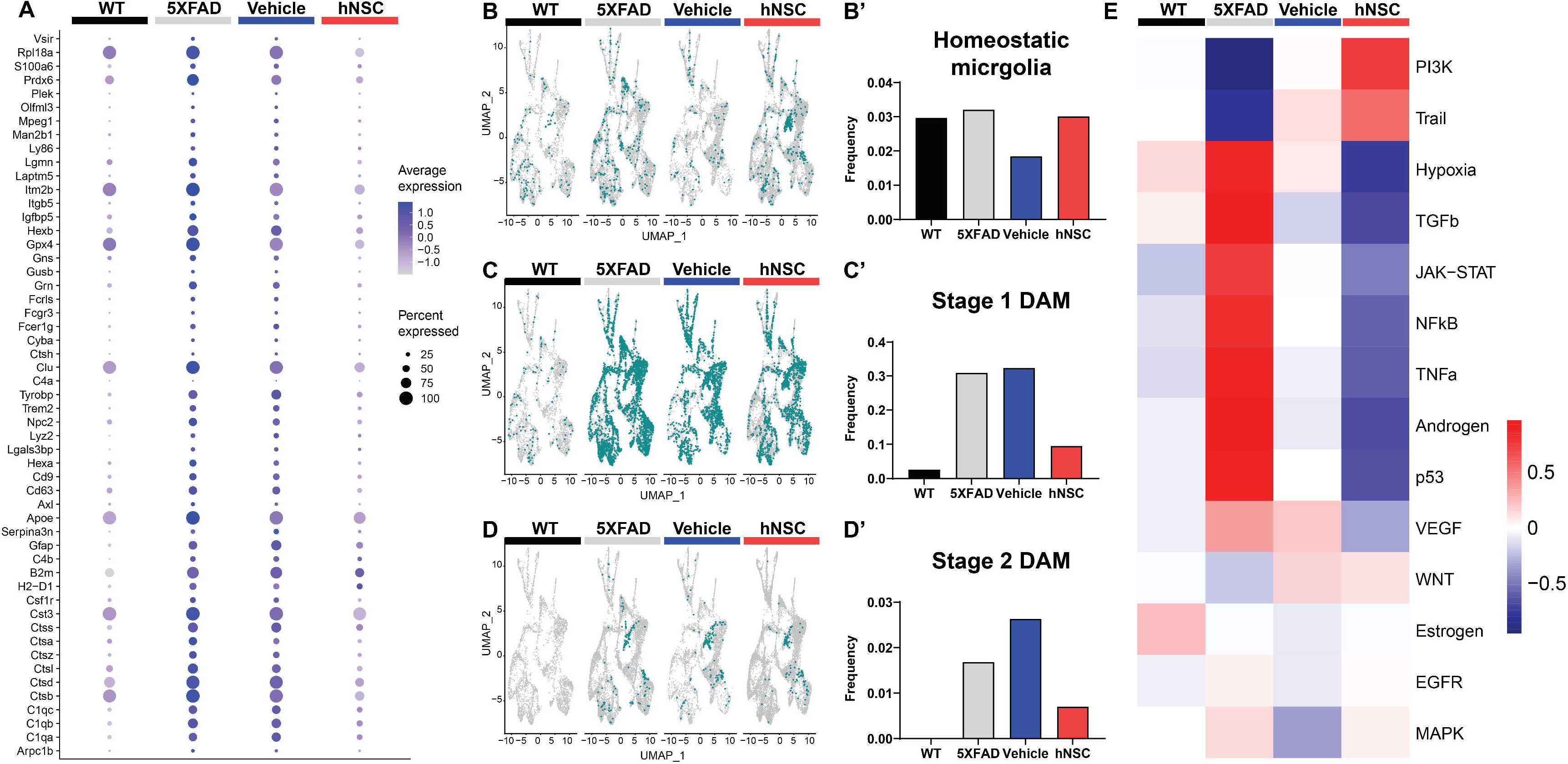
hNSC transplantation normalizes plaque-induced genes (PIGs) and microglial phenotypes in the brain of 5XFAD mice after 34 weeks. (**A**) PIGs feature plot in microglia for wild-type (WT, black), untreated 5XFAD (grey), immunosuppressed only (IS only) 5XFAD (blue), and hNSC treated 5XFAD (red) groups. Plot represents 57 PIGs (y-axis) for various groups (x-axis). PIGs average expression level represented by color scale from less (grey) to higher (purple) expression. Percent expressed represented by dot size from smallest (expressed by none) to largest (100%, expressed by all cells from the cluster). (**B-D**) UMAP plots for WT (black), untreated 5XFAD (grey), IS only 5XFAD (blue), and hNSC injected 5XFAD (red) for (**B**) homeostatic microglia (HMG), (**C**) stage 1 disease-associated microglia (DAMs), and (**D**) stage 2 DAMs represented as green dots. (**B’-D’**) Quantitation of (**B’**) HMG, (**C’**) stage 1 DAMs, and (**D’**) stage 2 DAMs as a proportion of total microglia for WT (black), untreated 5XFAD (grey), IS only 5XFAD (blue), and hNSC injected 5XFAD (red) groups. (**E**) Heatmap from PROGENy analysis for WT (black), untreated 5XFAD (grey), IS only 5XFAD (blue), and hNSC injected 5XFAD (red) groups. PROGENy scores represented by color scale from less (blue) to higher (red) scores.

Since microglia are so deeply connected to amyloid levels and response, we next investigated microglial phenotypes among the distinct conditions. We considered three phenotypes, non-pathological homeostatic microglia along with two states associated with a response to a pathologic condition, the stage 1 disease-associated microglia (DAMs) and stage 2 DAMs. DAMs are a microglial subset characterized by downregulated homeostatic genes (*e.g*., *Tmem119*, *Cx3cr1*, *P2ry12*) and various upregulated genes (*e.g*., *Trem2*, *ApoE*, inflammatory genes) (**Figure S11**) (Keren-Shaul et al., 2017; Chen and Colonna, 2021). DAMs progress from a stage 1 to a stage 2 category and are linked with the neuroimmune response to AD. The level of homeostatic microglia was relatively constant across WT, 5XFAD, IS only, and hNSC conditions (**Figures 5B, 5B’**). As anticipated, there were relatively few stage 1 DAMs in WT brain, which increased in frequency in 5XFAD brain (**Figures 5C, 5C’**). hNSC transplantation, but not IS only, lowered the number of brain stage 1 DAMs. Interestingly, no stage 2 DAMs were detected in WT brain, while multiple stage 2 DAMs were detected in 5XFAD animals, which were lowered by hNSC, but not IS only, treatment (**Figures 5D, 5D’**). Thus, hNSCs lowered the number of stage 1 and stage 2 DAMs associated with the pathologic AD state.

To gain further insight into inflammatory processes, we leveraged PROGENy analysis, which estimates the activity of relevant signaling pathways based on consensus gene signatures generated from a database of perturbation experiments, *i.e*., inflammatory activation (Schubert et al., 2018). PROGENy differs from conventional pathway enrichment methods by considering the effects of post-translational modifications and downstream signaling on inflammatory pathway activation. As anticipated, the brain of 5XFAD mice exhibited an inflammatory phenotype, with general activation of several inflammatory pathways, *e.g*., TNF-α, JAK/STAT (**Figure 5E**). IS only containing an immunosuppressive regimen lowered brain inflammation, as would be expected. However, hNSC transplantation exerted a stronger anti-inflammatory effect, suppressing TNFα, JAK/STAT, and NF-κB signaling pathways.

Overall, hNSC transplantation into 5XFAD mice reversed the expression of PIGs, particularly in microglia, and reduced the population of DAM phenotypes.

### hNSC transplantation alters cell signaling in the brain of 5XFAD mice after 34 weeks

Microglia constantly surveil the brain and respond to external signals (ElAli and Rivest, 2016); thus, cell-to-cell communication is an important aspect of microglial function. Moreover, transplanted stem cells exert beneficial effects via the “neighborhood” route, by enriching, and presumably communicating with, the local milieu. Thus, we investigated intercellular communication in the brain among all cell types, including microglia and hNSCs, using CellChat (Jin et al., 2021). This tool predicts intercellular communication by leveraging a database of more than 2,000 ligand-receptor pairs. CellChat presents overview data in information flow charts of intercellular signaling pathways by summing probabilities of communication averaged across all cell types. CellChat also presents more granular data in circle plots by homing in on interactions between specific cell types acting as signal sender (cells expresses signaling ligand) and signal receiver (cells expresses signaling receptor).

Overall, there were 2418, 4346, and 1017 inferred ligand-receptor interactions across cells of WT, 5XFAD, and hNSC conditions, respectively. Compared to WT brain cells, pathways that were significantly attenuated on average across all 5XFAD cells involved neuronal growth regulator (NEGR) and pleiotrophin (PTN), which play a significant role in neurite outgrowth among other functions (**Figure S12A**). Conversely, multiple signaling pathways were significantly enhanced on average across all 5XFAD versus WT communicating cells, including APP, adhesion (*e.g*., L1CAM, NCAM, LAMININ, COLLAGEN), angiogenesis (*e.g*., PECAM1, VEGF), mitogenesis (*e.g*., PDGF, FGF), neuronal growth cone guidance (*e.g*., SEMA3, SEMA4, SEMA5, SEMA6, SEMA7), and myelination (*e.g*., MPZ). The most significant KEGG pathways encompassing the most signaling interactions included “cell adhesion molecules”, “axon guidance”, and “PI3K−Akt signaling pathway” (**Figure S13A**; **Table S7**). Similarly, GO enrichment identified “cell adhesion” as most significant (**Figure S13B**; **Table S7**). When we examined signaling affected by stem cell transplantation, only one pathway was significantly enhanced across cells in hNSC-injected brain, which, was APP (**Figure S12B**). All other signaling continued to be enhanced in 5XFAD versus hNSC injected 5XFAD brain, related to “cell adhesion molecules” (KEGG; **Figure S13C**; **Table S8**) and “cell adhesion” and “nervous system development” along with additional similar pathways (GO; **Figure S13D**; **Table S8**).

### hNSC transplantation alters cell signaling from and to hNSCs in the brain of 5XFAD mice after 34 weeks

Examination by ligand-receptor signaling between specific cell types revealed extensive intercellular signaling networks between cells under all conditions (WT, 5XFAD, IS only, hNSCs) (**Figure S14; Tables S9-S10**). Next, we homed in on ligand-receptor interactions specifically from and to hNSCs. hNSCs signaled to all cell types except cortical cells (**Figure 6A**). Importantly, most signals from hNSCs were directed at stage 1 and stage 2 DAMs with 19 and 29 interactions, respectively. These ligand-receptor pairs spanned APP-CD74, PSAP-GPR37, and GRN-SORT1, among others (**Table S11**). Biological inference by KEGG was highly significant for hNSCs communicating through “cell adhesion molecules” signaling with both stage 1 and stage 2 DAMs and “axon guidance” signaling with stage 2 DAMs (**Figure S15**; **Table S12**), presumably countering these pathways in 5XFAD brain. GO pathway for hNSC signaling with stage 1 and stage 2 DAMs included “cell adhesion” along with several nervous system and neuronal genesis, development, and differentiation pathways, again counteracting this signaling in 5XFAD.

**Figure 6.**
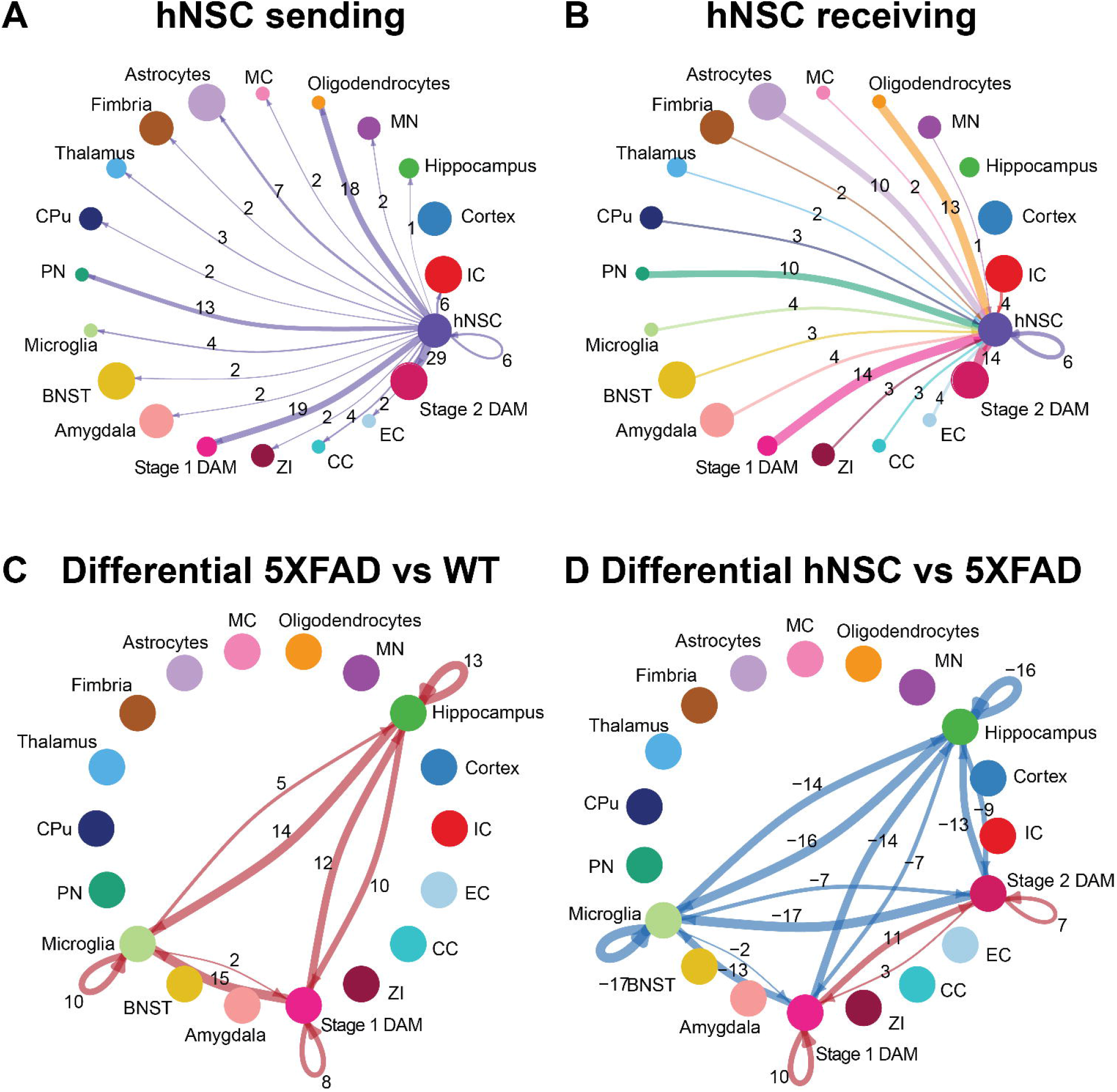
hNSC transplantation alters microglial cell signaling in the brain of 5XFAD mice after 34 weeks. Circle plots of ligand-receptor pair interactions between cell types with hNSCs as (**A**) sender and (**B**) receiver in the brain of hNSC injected 5XFAD mice. Dots represent distinct cell populations; strokes represent communication between hNSCs with distinct cell populations; loops represent hNSCs self-communication. In the circle plot of hNSCs as senders (**A**), stroke and loop colors are purple to represent signals originating from hNSCs. In the circle plot of hNSCs as receivers (**B**), stroke colors represent signals originating from distinct cell populations. Stroke and loop thickness reflects strength of the signaling pair, number reflects the number of ligand-receptor pair interactions. Circle plots of differential ligand-receptor pair interactions across cell types in (**C**) wild-type (WT) versus untreated 5XFAD and (**D**) hNSC injected versus untreated 5XFAD brain. Dots represent distinct cell populations; strokes represent communication between distinct cell populations; loops represent signaling within cell populations. Stroke and loop colors reflect upregulated (red) and downregulated (blue) differential ligand-receptor pair interactions. Stroke and loop thickness reflects strength of the differential signaling pair, number reflects the number of differential ligand-receptor pair interactions. BNST, bed nucleus of the stria terminalis; CC, corpus callosum; CPu, caudate putamen; DAM, disease associated microglia; EC, ependymal cells; IC, internal capsule; MC, meningeal cells; MN, medial nucleus; PN, piriform nucleus; ZI, zona incerta.

hNSCs also received signals from most cell types, except for hippocampal and cortical cells (**Figure 6B**). Again, hNSCs were the recipients of most signals from stage 1 and stage 2 DAMs, with 14 ligand-receptor interactions each, which once more encompassed APP-CD74, PSAP-GPR37, and GRN-SORT1, among others (**Table S13**). KEGG and GO pathways of these ligand-receptor interactions for stage 1 and stage 2 DAMs signaling to hNSCs were broadly reciprocated by pathway analysis of hNSC to stage 1 and stage 2 DAM signaling (*data not shown*).

### hNSC transplantation alters microglial cell signaling in the brain of 5XFAD mice after 34 weeks

Finally, we assessed differential ligand-receptor interactions among the various conditions across the hippocampus, microglia, and DAMs. 5XFAD increased the number of interactions between hippocampus, microglia, and stage 1 DAMs versus WT brain (positive numbers over strokes in **Figure 6C**; there are no stage 2 DAMs in WT, so this population is not present in the circle plot). These interactions were most significant and/or of higher probability for communication with PSAP-GPR37L1, which was downregulated in microglia signaling to hippocampus and other microglia in WT brain but upregulated between microglia and DAMs in 5XFAD brain (**Figure S16**). hNSC transplantation decreased the number of interactions between hippocampus and microglia to stage 1 and stage 2 DAMs versus 5XFAD brain (negative numbers over strokes in **Figure 6D**), countering AD pathology. hNSC regulated brain inflammation by either downregulating interaction pathways between microglia, stage 1 DAMs, and stage 2 DAMs (**Fig S16C**), while upregulating other interaction pathways (**Figure S16D**). Interestingly, the downregulated pathways by hNSC are the same pathways that were upregulated in 5XFAD versus WT (**Figure S12**). hNSC also decreased the activation of inflammatory pathways, including IGF, IL34, Thy1, SPP, and CX3CL1. Moreover, hNSCs upregulated communication between stage 1 and stage 2 DAMs, centered on APP, CADM, VTN, and PSAP signaling.

Overall, hNSC transplantation in 5XFAD hippocampus profoundly influenced communication across brain cells, especially with microglia and stage 1 and stage 2 DAMs.

## DISCUSSION

hNSC transplantation improved spatial memory in 5XFAD mice, without associated changes to amyloid plaque or cytokine levels. Spatial transcriptomics revealed amyloid-independent mechanisms of hNSCs, normalizing many DEGs in the brain and hippocampus. Treatment also restored PIGs and microglial phenotypes, along with ameliorating hippocampal/microglial signaling. Overall, hNSCs improved multiple aspects of AD, including cognition and underlying transcriptomic and microglial signatures.

Stem cells may treat neurodegenerative diseases, including AD (Boese et al., 2020), through multifactorial effects (Goutman et al., 2019; Sakowski and Chen, 2022). hNSC-mediated cognitive improvement has been confirmed by our group and others in many AD models (Ager et al., 2015; Kim et al., 2015; Li et al., 2016; McGinley et al., 2018). Additionally, mesenchymal stem cells (Matchynski-Franks et al., 2016) and induced pluripotent stem cells (Cha et al., 2017) ameliorate cognitive deficits in 5XFAD mice. Still, therapeutic mechanisms of stem cells in prior work remains ill-defined.

In the 5XFAD model, we found that hNSCs did not affect Aβ plaque burden. While some studies support cognitive improvement without change in amyloid burden (Ager et al., 2015; Li et al., 2016; Zhang et al., 2016), others report cognitive improvement linked to a decrease in Aβ plaques (Kim et al., 2015; Lee et al., 2015; McGinley et al., 2018). These disparate findings may arise from model-specific differences, age at transplantation, treatment duration, or the nature of stem cells used. Nevertheless, ample literature suggests hNSCs can improve cognition *via* mechanisms beyond amyloid. Inflammatory cytokine signaling is one oft-investigated pathway. Here, we observed that hNSCs did not impact cytokine levels on the whole-hemisphere level; however, we did identify effects on various cell adhesion proteins, growth factors, and extracellular matrix remodeling proteins. We previously noted similar results in IL-1β and TNF-α levels (McGinley et al., 2018), while other groups have shown hNSC-mediated suppression of IL-1β and TNF-α upon transplantation concomitant with changes in microglia activation state (Kim et al., 2015; Lee et al., 2015; Zhang et al., 2016).

Since hNSC mechanisms are likely to act through local signaling, we conducted a spatial transcriptomic analysis to assess hNSC-mediated “neighborhood” interactions beyond Aβ plaques and cytokines. Globally across the brain, there were 109 DEGs uniquely normalized in response to hNSCs (*i.e*., not downregulated by IS), several known to be key in AD risk and pathogenesis, such as *Trem2* (Keren-Shaul et al., 2017; Scheltens et al., 2021), *Tyrobp* (Pottier et al., 2016), *S100a1* (Afanador et al., 2014; Cristóvão and Gomes, 2019), *Igfbp5* (Barucker et al., 2015; Yu et al., 2018), *Gfap* (Benedet et al., 2021), and several complement genes (*C1qa*, *C1qb*, *C1qc*). Mutations to *Trem2* (Scheltens et al., 2021) and its adaptor protein *Tyrobp* (Pottier et al., 2016) are associated with increased AD risk. Elevated *Trem2* and *Tyrobp* are characteristic of DAMs, which aggregate around and phagocytose Aβ plaques as part of the neuroimmune response in AD (Keren-Shaul et al., 2017). hNSCs also affected S100A1, a regulator of tau phosphorylation, APP expression, and neuronal sensitivity to Aβ. S100A1 knockout in an AD mouse model reduced astrocytosis and microgliosis (Afanador et al., 2014; Cristóvão and Gomes, 2019). hNSC transplantation also lowered brain *Gfap* levels, which positively correlate with AD (Benedet et al., 2021). Another interesting gene reversed by hNSCs was insulin-like growth factor binding protein 5 (*Igfbp5*). IGFBP5 is elevated in AD mice (Barucker et al., 2015) and higher levels associate with faster cognitive decline in AD patients (Yu et al., 2018). Thus, the potential for hNSCs to impact multiple pathogenic signaling pathways is underscored here.

Next, we spatially restricted our analysis to the hippocampus, both as the center of memory consolidation adversely affected in AD, and as the brain structure most associated with the hNSC transplant target. There were far more DEGs in hippocampus (1061 uniquely corrected by hNSC transplant) than in whole brain analysis, where region-specific heterogeneity in DEGs might be averaged out. A number of DEGs in hippocampus are identifiable as AD-related genes, such as the transcription factor *Neurod6* regulating nervous system development and differentiation (Li et al., 2015), the peptidase *Ctsd* controlling protein turnover (Schuur et al., 2011), and glutaminase *Gls* (Griffin et al., 2018). Impaired cofilin (*Cfl1*) dynamics are also key in AD (Wang et al., 2020). In KEGG pathway enrichment analysis, ubiquitin mediated proteolysis was most significant and held the most DEGs, followed closely by protein processing in endoplasmic reticulum. Perturbations of both these pathways underlie Alzheimer’s pathophysiology (Hegde et al., 2019; Ghemrawi and Khair, 2020). Additional represented pathways included protein export, proteosome, and various RNA processing and metabolic pathways. GO pathway analysis identified cellular localization as the most significant and cellular macromolecule metabolic process as encompassing most DEGs. In addition, protein (macromolecule) and metabolic modification and localization also emerged in GO analysis.

In addition to our examination of all DEGs, we also focused our attention specifically on PIGs (Chen et al., 2020). Although hNSC transplantation did not directly affect global amyloid burden, we reasoned they may instead modulate the host response to Aβ plaques. Indeed, using our granular spatial dataset, we found that hNSC transplantation reversed 5XFAD-associated upregulation of PIGs, particularly in microglia. This primarily microglial signature of PIGs is confirmed by the literature (Chen et al., 2020). Moreover, PIGs overlap strongly with DAM genes; thus, we next examined the levels of homeostatic microglia and stage 1 and stage 2 DAMs. DAMs rise in numbers with AD progression: stage 1 DAMs represent an initial attempt to clear Aβ plaques (Keren-Shaul et al., 2017), a response which ultimately fails as chronically activated microglia adopt a stage 2 DAM phenotype later in disease (Heneka et al., 2015). Here, hNSC transplantation suppressed both stage 1 and stage 2 DAMs. This effect was not due to a reduction in total microglial populations, as the proportion of homeostatic microglia remained unchanged. One interpretation is that transplantation blunts the toxic downstream effects of amyloid plaques, obviating the need for DAMs. Anti-inflammatory characteristics of stem cells were replicated in the PROGENy analysis (Schubert et al., 2018), which found that hNSC transplantation opposed inflammation in the 5XFAD brain, by suppressing TNFα, JAK/STAT, and NF-κB signaling pathways.

Finally, we examined intercellular signaling and the potential positive benefits of hNSCs *via* local “neighborhood” interactions (Goutman et al., 2019). There was greatest intercellular communication in the 5XFAD brain, which was reduced by hNSCs. Enhanced communication between 5XFAD cells was mainly centered around cell adhesion and nervous system development. By contrast cells in hNSC transplanted brain communicated most notably through APP signaling. Although this may seem paradoxical, GO pathway enrichment analysis frequently placed APP signaling into various neurodevelopmental pathways, aligned with its roles in neuronal development, signaling, and intracellular transport in health (van der Kant and Goldstein, 2015); therefore, APP signaling in this context may represent a restorative process. Moreover, one study suggests that APP-CD74 interaction (the ligand-receptor pair for APP signaling), reduces Aβ production in a cell model (Matsuda et al., 2009), CD74 being the invariant chain of the class II major histocompatibility complex.

We next looked more closely at communication between specific cell types, and hNSCs particularly. hNSCs interacted the most with stage 1 and stage 2 DAMs. Some of the ligand-receptor interactions involved in hNSC-DAM communication, such as APP-CD74 and GRN-SORT1, are high or moderate risk genes for AD (Scheltens et al., 2021; Bellenguez et al., 2022). Finally, we completed our analysis by examining differential interactions between hippocampus, microglia, and DAMs in untreated and hNSC-transplanted 5XFAD brain. Again, interactions were reduced akin to wild type animals. Importantly, the most upregulated and significant signaling pathway between microglia and DAM communication in 5XFAD brain was PSAP-GPR37L1, related to lysosomal regulation, which accumulates in dystrophic neurites and/or Aβ plaques of AD mice (Sharoar et al., 2021) and patients (Mendsaikhan et al., 2019). hNSC transplantation reversed this signaling interaction. Additionally, hNSCs suppressed 5XFAD-induced inflammation, *e.g*., IL34, Thy1, SPP, and CX3CL1, indicative of an anti-inflammatory effect, as verified by other methods (*e.g*., DAM levels and PROGENy analysis).

This is the first study, to our knowledge, to implement spatial transcriptomic data to examine the long-term effects of hNSC transplantation. This facilitated analysis of region- and cell-specific changes, especially of the hippocampus, of PIGs in microglia, and of DAMs. Our bioinformatics analysis of transcriptional changes in the 5XFAD brain, and remediation by hNSCs, spanned multiple methods, including DEGs analysis, DAM levels, PROGENy analysis, and intercellular communication. However, our study also had limitations. First, behavioral and spatial transcriptomic cohorts were on different immunosuppressive regimens. Given that the tissues utilized for spatial transcriptomics aimed to investigate longer-term hNSC effects, we employed an immunosuppressive regimen developed specifically to enhance longer-term graft survival (McGinley et al., 2022). Moreover, the need for immunosuppression is a study weakness, albeit a necessary one when using a xenograft model of human stem cells in immune competent mice. An alternative strategy is to use immunodeficient mice; however, since immune cells, and microglia especially (Heneka et al., 2015), are integral to AD pathology, we feel such a model would not have faithfully reproduced the full spectrum of hNSC mechanisms. We therefore opted to use immune competent mice, and our analysis utilizes an immunosuppression-alone cohort to exclude any spurious findings secondary to immunosuppression. Second, we applied only one well-established behavioral test, the Morris Water Maze, but confirmatory experiments using additional hippocampal-dependent and cognitive tests are warranted. Third, we only employed male mice, although sex differences are anticipated in the 5XFAD model (Sil et al., 2022) and in the pathology discussed, such as neuroinflammation and microglial biology (Sala Frigerio et al., 2019).

## CONCLUSIONS

Overall, our findings cumulatively suggest that stem cells exert pleiotropic salutary effects, clarifying the mechanisms of this promising therapeutic strategy. Stem cell transplantation into the brain of 5XFAD mice rescues cognitive deficits, and spatial transcriptomics suggests improvements in protein processing pathways to be a major contributor. Additionally, hNSCs interacted with and normalized populations of stage 1 and stage 2 DAMs and suppressed PIG expression and inflammatory signaling despite stable Aβ plaque levels. Advances in AD treatment may require multi-pronged approaches beyond amyloid, and our findings here suggest that a paradigm incorporating microglial-modulating effects may offer unique therapeutic opportunities.

## Authors’ contributions

Conceptualization KSC, LMG, ELF; Methodology KSC, LMG, MHN, ELF; Formal analysis KSC, MHN, JFK; Software MHN; Investigation LMG, JMH, DMR, JFK, SNM, FEM; Writing - Original Draft KSC, MHN, MGS, ELF; Writing - Review & Editing all authors; Visualization MHN, MGS; Supervision KSC, ELF; Project administration ELF; Funding acquisition KSC, LMG, ELF.

## Acknowledgements

The authors thank Palisade Bio, Inc. for supplying the hNSC line used in the study. They would like to acknowledge the University of Michigan Advanced Genomics Core.

## Data Availability Statement

All data supporting the findings of this study are available within the paper and its supplementary materials. The spatial transcriptomics data from this publication have been deposited in NCBI Gene Expression Omnibus (16) and are accessible through GEO Series accession number GSE209583 (https://www.ncbi.nlm.nih.gov/geo/query/acc.cgi?acc=GSE209583 and enter token gzglogqkvjqrhmt). Additional supporting data are available from the corresponding author upon reasonable request.

## REFERENCES

1. Afanador, L., Roltsch, E.A., Holcomb, L., Campbell, K.S., Keeling, D.A., Zhang, Y., et al. (2014). The Ca2+ sensor S100A1 modulates neuroinflammation, histopathology and Akt activity in the PSAPP Alzheimer’s disease mouse model. Cell Calcium 56(2), 68–80. doi: 10.1016/j.ceca.2014.05.002.

2. Ager, R.R., Davis, J.L., Agazaryan, A., Benavente, F., Poon, W.W., LaFerla, F.M., et al. (2015). Human neural stem cells improve cognition and promote synaptic growth in two complementary transgenic models of Alzheimer’s disease and neuronal loss. Hippocampus 25(7), 813–826. doi: 10.1002/hipo.22405.

3. Alves, R., and Higdon, R. (2013). “Differential Expression Analysis,” in Encyclopedia of Systems Biology, eds. W. Dubitzky, O. Wolkenhauer, K.-H. Cho & H. Yokota. (New York, NY: Springer New York), 572–572.

4. Arranz, A.M., and De Strooper, B. (2019). The role of astroglia in Alzheimer’s disease: pathophysiology and clinical implications. Lancet Neurol 18(4), 406–414. doi: 10.1016/s1474-4422(18)30490-3.

5. Barucker, C., Sommer, A., Beckmann, G., Eravci, M., Harmeier, A., Schipke, C.G., et al. (2015). Alzheimer amyloid peptide aβ42 regulates gene expression of transcription and growth factors. J Alzheimers Dis 44(2), 613–624. doi: 10.3233/jad-141902.

6. Bellenguez, C., Küçükali, F., Jansen, I.E., Kleineidam, L., Moreno-Grau, S., Amin, N., et al. (2022). New insights into the genetic etiology of Alzheimer’s disease and related dementias. Nature Genetics 54(4), 412–436. doi: 10.1038/s41588-022-01024-z.

7. Benedet, A.L., Milà-Alomà, M., Vrillon, A., Ashton, N.J., Pascoal, T.A., Lussier, F., et al. (2021). Differences Between Plasma and Cerebrospinal Fluid Glial Fibrillary Acidic Protein Levels Across the Alzheimer Disease Continuum. JAMA Neurol 78(12), 1471–1483. doi: 10.1001/jamaneurol.2021.3671.

8. Boese, A.C., Hamblin, M.H., and Lee, J.P. (2020). Neural stem cell therapy for neurovascular injury in Alzheimer’s disease. Exp Neurol 324, 113112. doi: 10.1016/j.expneurol.2019.113112.

9. Brettschneider, J., Del Tredici, K., Lee, V.M., and Trojanowski, J.Q. (2015). Spreading of pathology in neurodegenerative diseases: a focus on human studies. Nat Rev Neurosci 16(2), 109–120. doi: 10.1038/nrn3887.

10. Bromley-Brits, K., Deng, Y., and Song, W. (2011). Morris water maze test for learning and memory deficits in Alzheimer’s disease model mice. J Vis Exp (53). doi: 10.3791/2920.

11. Butler, A., Hoffman, P., Smibert, P., Papalexi, E., and Satija, R. (2018). Integrating single-cell transcriptomic data across different conditions, technologies, and species. Nat Biotechnol 36(5), 411–420. doi: 10.1038/nbt.4096.

12. Cha, M.Y., Kwon, Y.W., Ahn, H.S., Jeong, H., Lee, Y.Y., Moon, M., et al. (2017). Protein-Induced Pluripotent Stem Cells Ameliorate Cognitive Dysfunction and Reduce Aβ Deposition in a Mouse Model of Alzheimer’s Disease. Stem Cells Transl Med 6(1), 293–305. doi: 10.5966/sctm.2016-0081.

13. Chen, W.T., Lu, A., Craessaerts, K., Pavie, B., Sala Frigerio, C., Corthout, N., et al. (2020). Spatial Transcriptomics and In Situ Sequencing to Study Alzheimer’s Disease. Cell 182(4), 976–991.e919. doi: 10.1016/j.cell.2020.06.038.

14. Chen, Y., and Colonna, M. (2021). Microglia in Alzheimer’s disease at single-cell level. Are there common patterns in humans and mice? J Exp Med 218(9). doi: 10.1084/jem.20202717.

15. Choudhary, S., and Satija, R. (2022). Comparison and evaluation of statistical error models for scRNA-seq. Genome Biol 23(1), 27. doi: 10.1186/s13059-021-02584-9.

16. Cristóvão, J.S., and Gomes, C.M. (2019). S100 Proteins in Alzheimer’s Disease. Front Neurosci 13, 463. doi: 10.3389/fnins.2019.00463.

17. Deczkowska, A., Keren-Shaul, H., Weiner, A., Colonna, M., Schwartz, M., and Amit, I. (2018). Disease-Associated Microglia: A Universal Immune Sensor of Neurodegeneration. Cell 173(5), 1073–1081. doi: 10.1016/j.cell.2018.05.003.

18. ElAli, A., and Rivest, S. (2016). Microglia Ontology and Signaling. Front Cell Dev Biol 4, 72. doi: 10.3389/fcell.2016.00072.

19. Garden, G.A., and Campbell, B.M. (2016). Glial biomarkers in human central nervous system disease. Glia 64(10), 1755–1771. doi: 10.1002/glia.22998.

20. Ghemrawi, R., and Khair, M. (2020). Endoplasmic Reticulum Stress and Unfolded Protein Response in Neurodegenerative Diseases. Int J Mol Sci 21(17). doi: 10.3390/ijms21176127.

21. Goutman, S.A., Savelieff, M.G., Sakowski, S.A., and Feldman, E.L. (2019). Stem cell treatments for amyotrophic lateral sclerosis: a critical overview of early phase trials. Expert Opin Investig Drugs 28(6), 525–543. doi: 10.1080/13543784.2019.1627324.

22. Griffin, J.W.D., Liu, Y., Bradshaw, P.C., and Wang, K. (2018). In Silico Preliminary Association of Ammonia Metabolism Genes GLS, CPS1, and GLUL with Risk of Alzheimer’s Disease, Major Depressive Disorder, and Type 2 Diabetes. J Mol Neurosci 64(3), 385–396. doi: 10.1007/s12031-018-1035-0.

23. Hafemeister, C., and Satija, R. (2019). Normalization and variance stabilization of single-cell RNA-seq data using regularized negative binomial regression. Genome Biol 20(1), 296. doi: 10.1186/s13059-019-1874-1.

24. Harris, M.A., Clark, J., Ireland, A., Lomax, J., Ashburner, M., Foulger, R., et al. (2004). The Gene Ontology (GO) database and informatics resource. Nucleic Acids Res 32(Database issue), D258–261. doi: 10.1093/nar/gkh036.

25. Hegde, A.N., Smith, S.G., Duke, L.M., Pourquoi, A., and Vaz, S. (2019). Perturbations of Ubiquitin-Proteasome-Mediated Proteolysis in Aging and Alzheimer’s Disease. Front Aging Neurosci 11, 324. doi: 10.3389/fnagi.2019.00324.

26. Heneka, M.T., Carson, M.J., El Khoury, J., Landreth, G.E., Brosseron, F., Feinstein, D.L., et al. (2015). Neuroinflammation in Alzheimer’s disease. Lancet Neurol 14(4), 388–405. doi: 10.1016/s1474-4422(15)70016-5.

27. Holland, C.H., Szalai, B., and Saez-Rodriguez, J. (2020). Transfer of regulatory knowledge from human to mouse for functional genomics analysis. Biochim Biophys Acta Gene Regul Mech 1863(6), 194431. doi: 10.1016/j.bbagrm.2019.194431.

28. Jin, S., Guerrero-Juarez, C.F., Zhang, L., Chang, I., Ramos, R., Kuan, C.H., et al. (2021). Inference and analysis of cell-cell communication using CellChat. Nat Commun 12(1), 1088. doi: 10.1038/s41467-021-21246-9.

29. Kanehisa, M., and Goto, S. (2000). KEGG: kyoto encyclopedia of genes and genomes. Nucleic Acids Res 28(1), 27–30. doi: 10.1093/nar/28.1.27.

30. Keren-Shaul, H., Spinrad, A., Weiner, A., Matcovitch-Natan, O., Dvir-Szternfeld, R., Ulland, T.K., et al. (2017). A Unique Microglia Type Associated with Restricting Development of Alzheimer’s Disease. Cell 169(7), 1276–1290.e1217. doi: 10.1016/j.cell.2017.05.018.

31. Kim, J.A., Ha, S., Shin, K.Y., Kim, S., Lee, K.J., Chong, Y.H., et al. (2015). Neural stem cell transplantation at critical period improves learning and memory through restoring synaptic impairment in Alzheimer’s disease mouse model. Cell Death Dis 6(6), e1789. doi: 10.1038/cddis.2015.138.

32. Lee, I.S., Jung, K., Kim, I.S., Lee, H., Kim, M., Yun, S., et al. (2015). Human neural stem cells alleviate Alzheimer-like pathology in a mouse model. Mol Neurodegener 10, 38. doi: 10.1186/s13024-015-0035-6.

33. Lein, E.S., Hawrylycz, M.J., Ao, N., Ayres, M., Bensinger, A., Bernard, A., et al. (2007). Genome-wide atlas of gene expression in the adult mouse brain. Nature 445(7124), 168–176. doi: 10.1038/nature05453.

34. Li, X., Long, J., He, T., Belshaw, R., and Scott, J. (2015). Integrated genomic approaches identify major pathways and upstream regulators in late onset Alzheimer’s disease. Sci Rep 5, 12393. doi: 10.1038/srep12393.

35. Li, X., Zhu, H., Sun, X., Zuo, F., Lei, J., Wang, Z., et al. (2016). Human Neural Stem Cell Transplantation Rescues Cognitive Defects in APP/PS1 Model of Alzheimer’s Disease by Enhancing Neuronal Connectivity and Metabolic Activity. Front Aging Neurosci 8, 282. doi: 10.3389/fnagi.2016.00282.

36. Matchynski-Franks, J.J., Pappas, C., Rossignol, J., Reinke, T., Fink, K., Crane, A., et al. (2016). Mesenchymal Stem Cells as Treatment for Behavioral Deficits and Neuropathology in the 5xFAD Mouse Model of Alzheimer’s Disease. Cell Transplant 25(4), 687–703. doi: 10.3727/096368916x690818.

37. Matsuda, S., Matsuda, Y., and D’Adamio, L. (2009). CD74 interacts with APP and suppresses the production of Abeta. Mol Neurodegener 4, 41. doi: 10.1186/1750-1326-4-41.

38. McGinley, L.M., Chen, K.S., Mason, S.N., Rigan, D.M., Kwentus, J.F., Hayes, J.M., et al. (2022). Monoclonal antibody-mediated immunosuppression enables long-term survival of transplanted human neural stem cells in mouse brain. Clin Transl Med 12(9), e1046. doi: 10.1002/ctm2.1046.

39. McGinley, L.M., Kashlan, O.N., Bruno, E.S., Chen, K.S., Hayes, J.M., Kashlan, S.R., et al. (2018). Human neural stem cell transplantation improves cognition in a murine model of Alzheimer’s disease. Sci Rep 8(1), 14776. doi: 10.1038/s41598-018-33017-6.

40. McGinley, L.M., Sims, E., Lunn, J.S., Kashlan, O.N., Chen, K.S., Bruno, E.S., et al. (2016). Human Cortical Neural Stem Cells Expressing Insulin-Like Growth Factor-I: A Novel Cellular Therapy for Alzheimer’s Disease. Stem Cells Transl Med 5(3), 379–391. doi: 10.5966/sctm.2015-0103.

41. Mendsaikhan, A., Tooyama, I., Bellier, J.P., Serrano, G.E., Sue, L.I., Lue, L.F., et al. (2019). Characterization of lysosomal proteins Progranulin and Prosaposin and their interactions in Alzheimer’s disease and aged brains: increased levels correlate with neuropathology. Acta Neuropathol Commun 7(1), 215. doi: 10.1186/s40478-019-0862-8.

42. Pottier, C., Ravenscroft, T.A., Brown, P.H., Finch, N.A., Baker, M., Parsons, M., et al. (2016). TYROBP genetic variants in early-onset Alzheimer’s disease. Neurobiol Aging 48, 222.e229–222.e215. doi: 10.1016/j.neurobiolaging.2016.07.028.

43. Sakowski, S.A., and Chen, K.S. (2022). Stem cell therapy for central nervous system disorders: Metabolic interactions between transplanted cells and local microenvironments. Neurobiol Dis 173, 105842. doi: 10.1016/j.nbd.2022.105842.

44. Sala Frigerio, C., Wolfs, L., Fattorelli, N., Thrupp, N., Voytyuk, I., Schmidt, I., et al. (2019). The Major Risk Factors for Alzheimer’s Disease: Age, Sex, and Genes Modulate the Microglia Response to Aβ Plaques. Cell Rep 27(4), 1293–1306.e1296. doi: 10.1016/j.celrep.2019.03.099.

45. Satija, R., Farrell, J.A., Gennert, D., Schier, A.F., and Regev, A. (2015). Spatial reconstruction of single-cell gene expression data. Nat Biotechnol 33(5), 495–502. doi: 10.1038/nbt.3192.

46. Savelieff, M.G., Lee, S., Liu, Y., and Lim, M.H. (2013). Untangling amyloid-β, tau, and metals in Alzheimer’s disease. ACS Chem Biol 8(5), 856–865. doi: 10.1021/cb400080f.

47. Scheltens, P., De Strooper, B., Kivipelto, M., Holstege, H., Chételat, G., Teunissen, C.E., et al. (2021). Alzheimer’s disease. Lancet 397(10284), 1577–1590. doi: 10.1016/s0140-6736(20)32205-4.

48. Schindelin, J., Rueden, C.T., Hiner, M.C., and Eliceiri, K.W. (2015). The ImageJ ecosystem: An open platform for biomedical image analysis. Mol Reprod Dev 82(7-8), 518–529. doi: 10.1002/mrd.22489.

49. Schubert, M., Klinger, B., Klünemann, M., Sieber, A., Uhlitz, F., Sauer, S., et al. (2018). Perturbation-response genes reveal signaling footprints in cancer gene expression. Nat Commun 9(1), 20. doi: 10.1038/s41467-017-02391-6.

50. Schuur, M., Ikram, M.A., van Swieten, J.C., Isaacs, A., Vergeer-Drop, J.M., Hofman, A., et al. (2011). Cathepsin D gene and the risk of Alzheimer’s disease: a population-based study and meta-analysis. Neurobiol Aging 32(9), 1607–1614. doi: 10.1016/j.neurobiolaging.2009.10.011.

51. Serrano-Pozo, A., Das, S., and Hyman, B.T. (2021). APOE and Alzheimer’s disease: advances in genetics, pathophysiology, and therapeutic approaches. Lancet Neurol 20(1), 68–80. doi: 10.1016/s1474-4422(20)30412-9.

52. Sharoar, M.G., Palko, S., Ge, Y., Saido, T.C., and Yan, R. (2021). Accumulation of saposin in dystrophic neurites is linked to impaired lysosomal functions in Alzheimer’s disease brains. Mol Neurodegener 16(1), 45. doi: 10.1186/s13024-021-00464-1.

53. Shi, J., Sabbagh, M.N., and Vellas, B. (2020). Alzheimer’s disease beyond amyloid: strategies for future therapeutic interventions. Bmj 371, m3684. doi: 10.1136/bmj.m3684.

54. Sil, A., Erfani, A., Lamb, N., Copland, R., Riedel, G., and Platt, B. (2022). Sex Differences in Behavior and Molecular Pathology in the 5XFAD Model. J Alzheimers Dis 85(2), 755–778. doi: 10.3233/jad-210523.

55. van der Kant, R., and Goldstein, L.S. (2015). Cellular functions of the amyloid precursor protein from development to dementia. Dev Cell 32(4), 502–515. doi: 10.1016/j.devcel.2015.01.022.

56. Vestal, B.E., Wynn, E., and Moore, C.M. (2022). lmerSeq: an R package for analyzing transformed RNA-Seq data with linear mixed effects models. BMC Bioinformatics 23(1), 489. doi: 10.1186/s12859-022-05019-9.

57. Wang, Q., Yuan, W., Yang, X., Wang, Y., Li, Y., and Qiao, H. (2020). Role of Cofilin in Alzheimer’s Disease. Front Cell Dev Biol 8, 584898. doi: 10.3389/fcell.2020.584898.

58. Yu, L., Petyuk, V.A., Gaiteri, C., Mostafavi, S., Young-Pearse, T., Shah, R.C., et al. (2018). Targeted brain proteomics uncover multiple pathways to Alzheimer’s dementia. Ann Neurol 84(1), 78–88. doi: 10.1002/ana.25266.

59. Zhang, Q., Wu, H.H., Wang, Y., Gu, G.J., Zhang, W., and Xia, R. (2016). Neural stem cell transplantation decreases neuroinflammation in a transgenic mouse model of Alzheimer’s disease. J Neurochem 136(4), 815–825. doi: 10.1111/jnc.13413.

