## Supplemental Figures and Table Legends for "Human neural stem cells restore spatial memory in a transgenic Alzheimer’s disease mouse model by an immunomodulating mechanism"

**Figure S1. hNSCs transplantation does not alter brain cytokine levels in 5XFAD after 8 weeks.** Multiplex ELISA of brain cytokines from hemibrain homogenates for (**A**) interleukins (IL), IL-1β, IL-2, IL-6, IL-10, and IL-12, (**B**) tumor necrosis factor α (TNF-α), (**C**) granulocyte-macrophage colony-stimulating factor (GM-CSF), (**D**) monocyte chemoattractant protein 1 (MCP-1), and (**E**) transforming growth factor (TGF), TGF-β1, TGF-β2, and TGF-β3, for wild-type (WT, black), untreated 5XFAD (grey), vehicle injected 5XFAD (blue), and hNSC injected 5XFAD (red) mice; n=5-9, *p<0.05, **p<0.01, ***p<0.001, by one-way ANOVA.

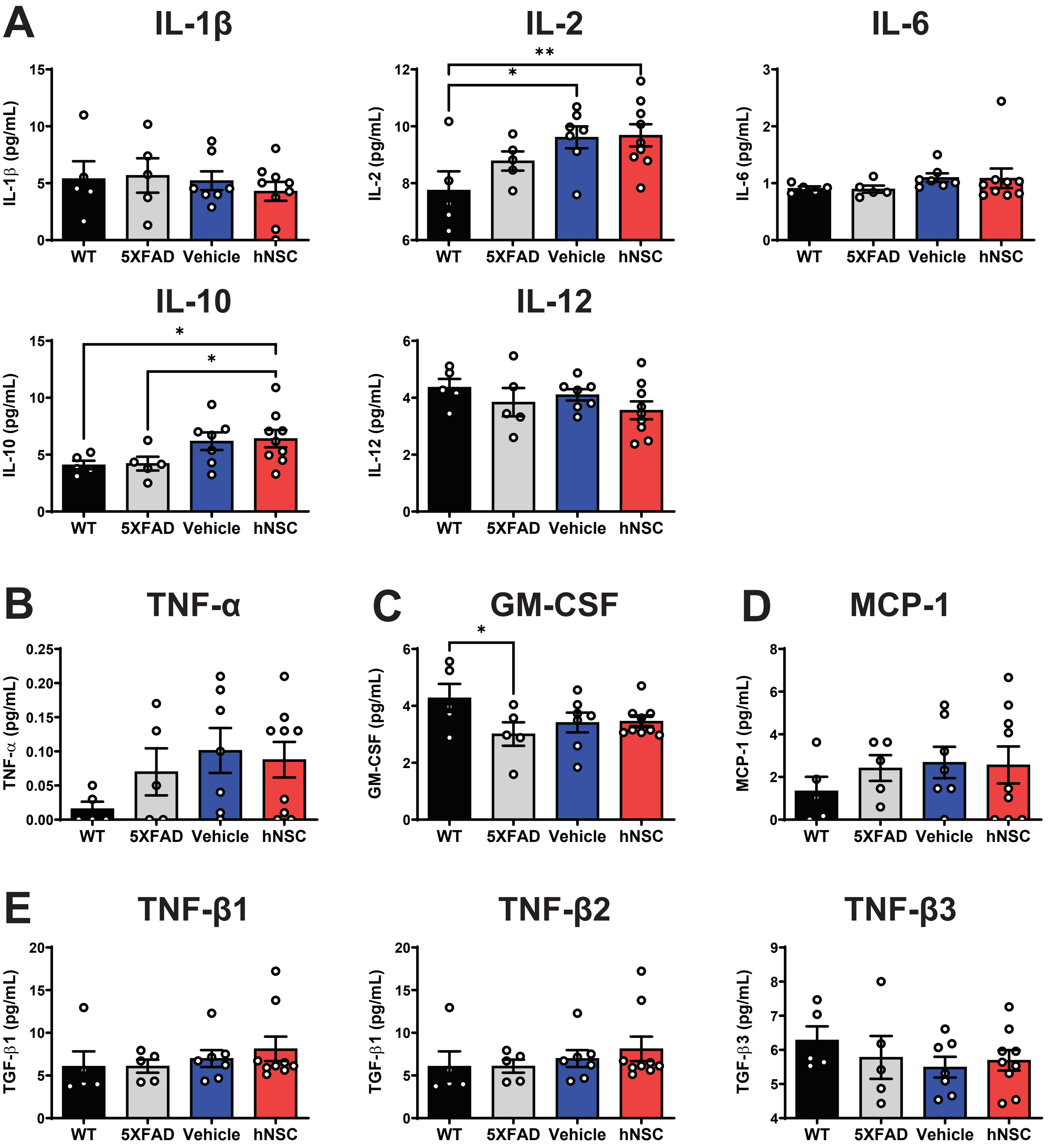

**Figure S2. hNSCs transplantation alters brain NCAM, sVCAM-1, and cathepsin D levels in 5XFAD after 8 weeks.** Multiplex ELISA of brain adhesion and extracellular matrix proteins from hemibrain homogenates for (**A**) neural cell adhesion molecule (NCAM), (**B**) soluble vascular cell adhesion protein 1 (sVCAM-1), (**C**) soluble intercellular adhesion molecule 1 (sICAM-1), (**D**) platelet-derived growth factor AA (PDGF-AA), (**E**) PDGF-AB, (**F**) cathepsin D, and (**G**) matrix metalloproteinase (MMP), MMP-2, MMP-3, MMP-8, MMP-9, and MMP-12, for wild-type (WT, black), untreated 5XFAD (grey), vehicle injected 5XFAD (blue), and hNSC injected 5XFAD (red) mice; n=5-9, *p<0.05, **p<0.01, ****p<0.0001, by one-way ANOVA.

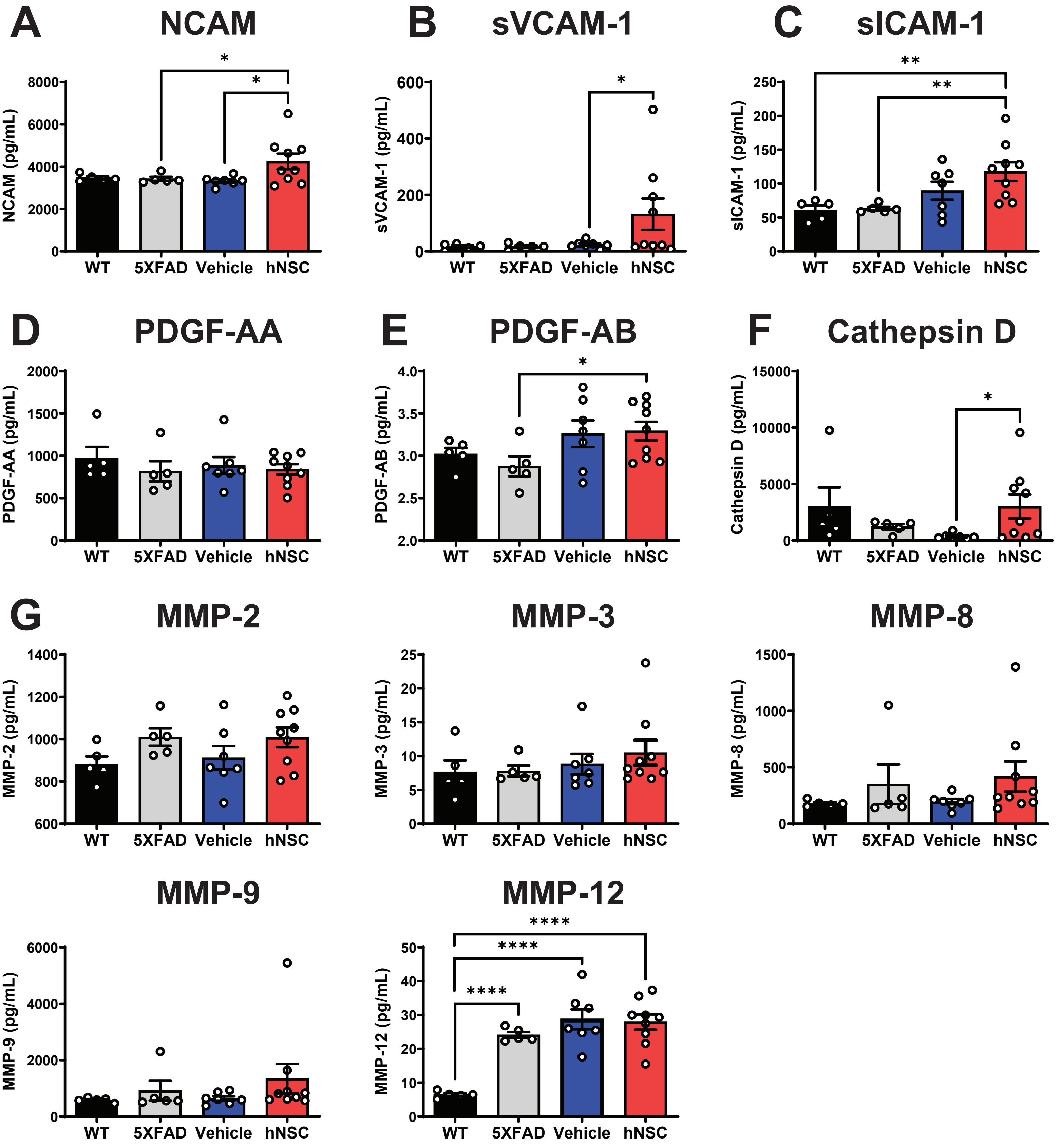

**Figure S3. hNSC transplantation does not alter brain Aβ levels in 5XFAD mice after 34 weeks.** Immunohistochemistry quantification by percent Aβ immunoreactivity area to total area in (**A**) dentate gyrus, (**B**) CA1, (**C**) CA3, and (**D**) fimbria fornix for hNSC injected 5XFAD (red), untreated 5XFAD (grey), wild-type (WT, black), and immunosuppression (IS) only (anti-CD4, -CD40L) 5XFAD (blue); *p<0.05, **p<0.01, ***p<0.001, by one-way ANOVA adjusted with Tukey's multiple comparisons.

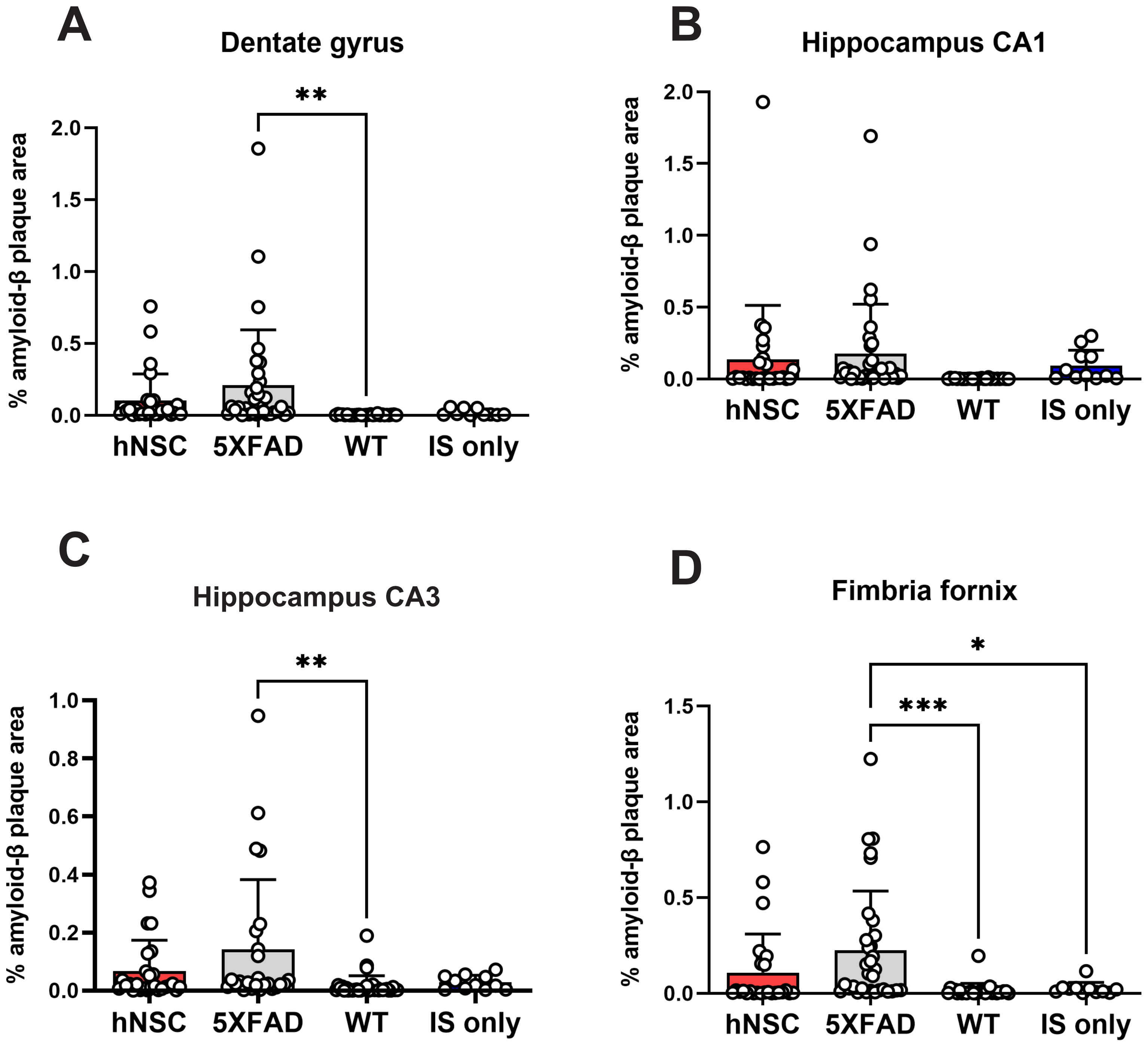

**Figure S4. Clustering of the spatial transcriptomics dataset maps to anatomic regions.** (**A**) Clustering of the spatial transcriptomics dataset by Uniform Manifold Approximation and Projection (UMAP) yields 25 cell populations (Garden and Campbell 2016, Deczkowska, Keren-Shaul et al. 2018). (**B-C**) Cell clusters map to their respective anatomic regions. (**D**) UMAP clustering of the spatial transcriptomics dataset yields 10 brain regions [Allen Brain Atlas (Lein, Hawrylycz et al. 2007)].
AN, amygdalar nucleus; BNST, bed nucleus of the stria terminalis; CA, ML, cornu ammonis, molecular layer; CA1, PL, cornu ammonis 1, pyramidal layer; CA3, PL, cornu ammonis 1, pyramidal layer; CC, corpus callosum; CP, choroid plexus; CPu, caudate putamen; DG, GL, dentate gyrus, granule cell layer; DG, ML, dentate gyrus, molecular layer; EC, ependymal cells; IC, internal capsule; L1, cortical layer 1; L2/3, cortical layer 2/3; L4, cortical layer 4; L5, cortical layer 5; L6a, L6b, cortical layers 6a and 6b; MC, meningeal cells; MG, microglia; MN, medial nucleus; PN, piriform nucleus; ZI, zona incerta.

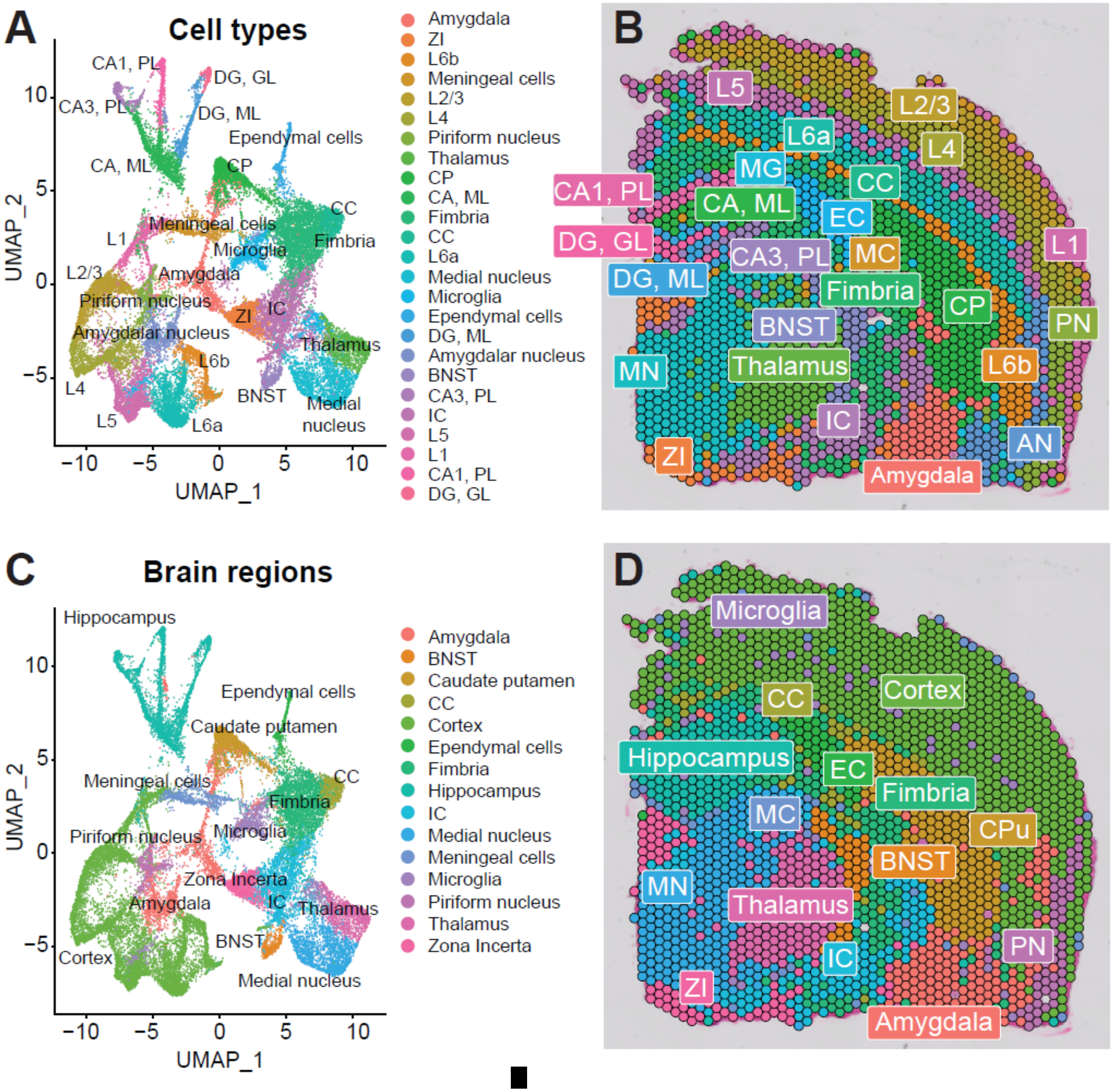

**Figure S5. hNSC gene expression.** Heatmap of the top 100 genes (x-axis) expressed by hNSCs. Y-axis represents cell barcodes for transplanted hNSCs. Abundance represented by color scale from less (blue) to more (red) abundant.

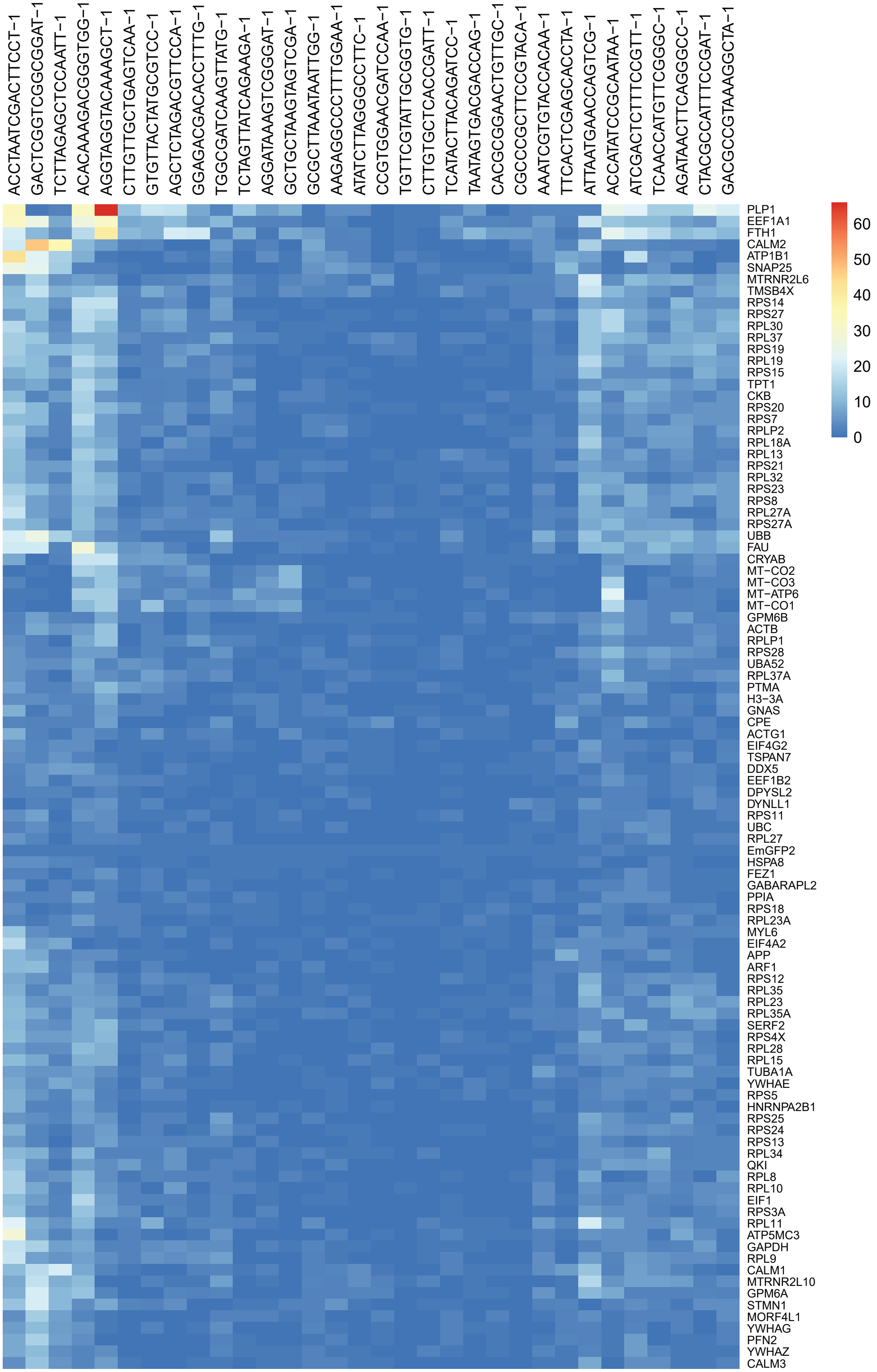

**Figure S6. hNSC pathway analysis.** Pathway analysis from top 100 hNSC genes by (**A**) Kyoto Encyclopedia of Genes and Genomes (KEGG) and (**B**) Gene Ontology (GO). Pathway significance represented by color scale from less (pink) to more (red) significant. Number over each bar represents the number of hNSC genes; rich factor represents the proportion of hNSC genes relative to the total number in the pathway.

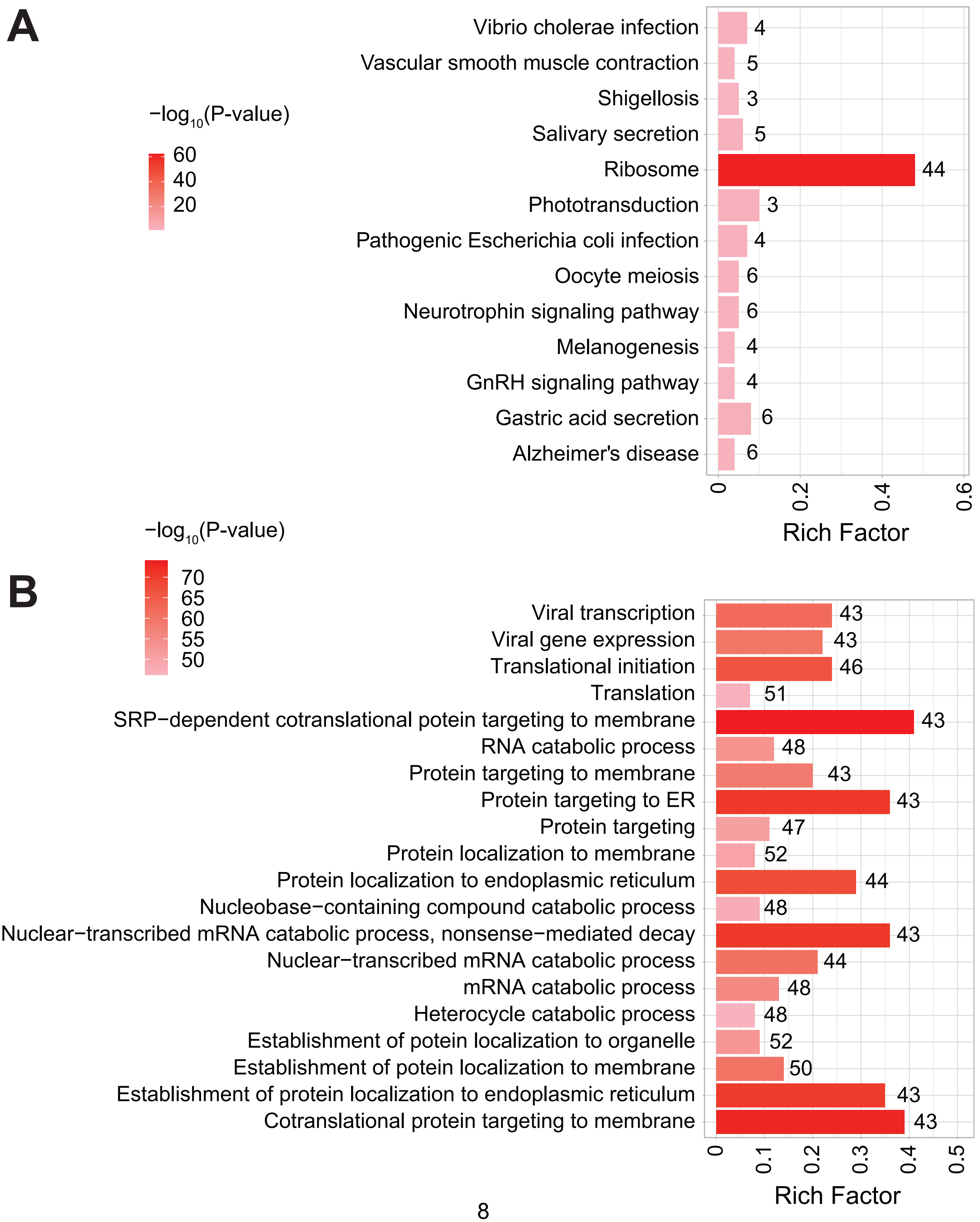

**Figure S7. hNSC transplantation normalizes differentially expressed genes (DEGs) in the brain of 5XFAD mice after 34 weeks; analysis of DEGs upregulated in 5XFAD versus WT and uniquely downregulated in hNSC versus 5XFAD.** (**A**) Venn diagram of upregulated DEGs in 5XFAD versus wild-type (WT; grey) and downregulated DEGs in immunosuppression only (IS only) versus 5XFAD (blue) and hNSC versus 5XFAD (red) in whole brain. (**B**) Bar plot of top 50 DEGs uniquely upregulated in 5XFAD versus WT (grey) and downregulated in hNSC versus 5XFAD (red) in whole brain. FC, fold-change. Entire DEGs list in Table S1. (**C**) KEGG (top panel) and GO (bottom panel) pathway analysis of 109 DEGs uniquely upregulated in 5XFAD versus WT and downregulated in hNSC versus 5XFAD in whole brain. Pathway significance represented by color scale from less (light green) to more (dark green) significant. Number over each bar represents the number of DEGs; rich factor represents the proportion of DEGs relative to the total number in the pathway.

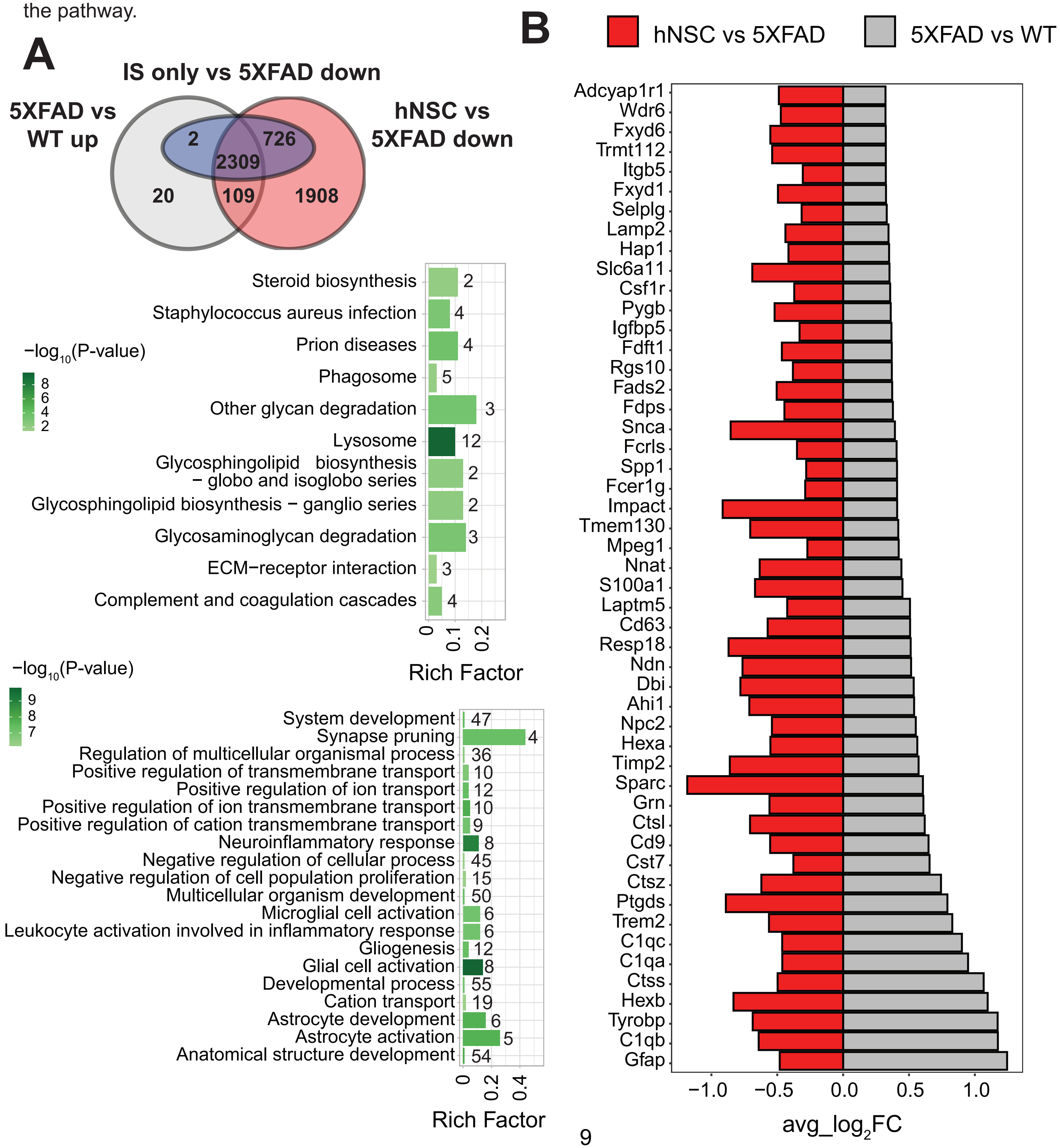

**Figure S8. Spatial and cluster distrbution of differentially expressed genes (DEGs).** (**A-B**) Represenative brains sections of Trem2 DEG across wild-type (WT, black), 5XFAD (grey), immunosuppression only (IS only; blue), and hNSC (red) brain, represented (**A**) spatially and (**B**) by UMAP plot; expression level indicated by scale.

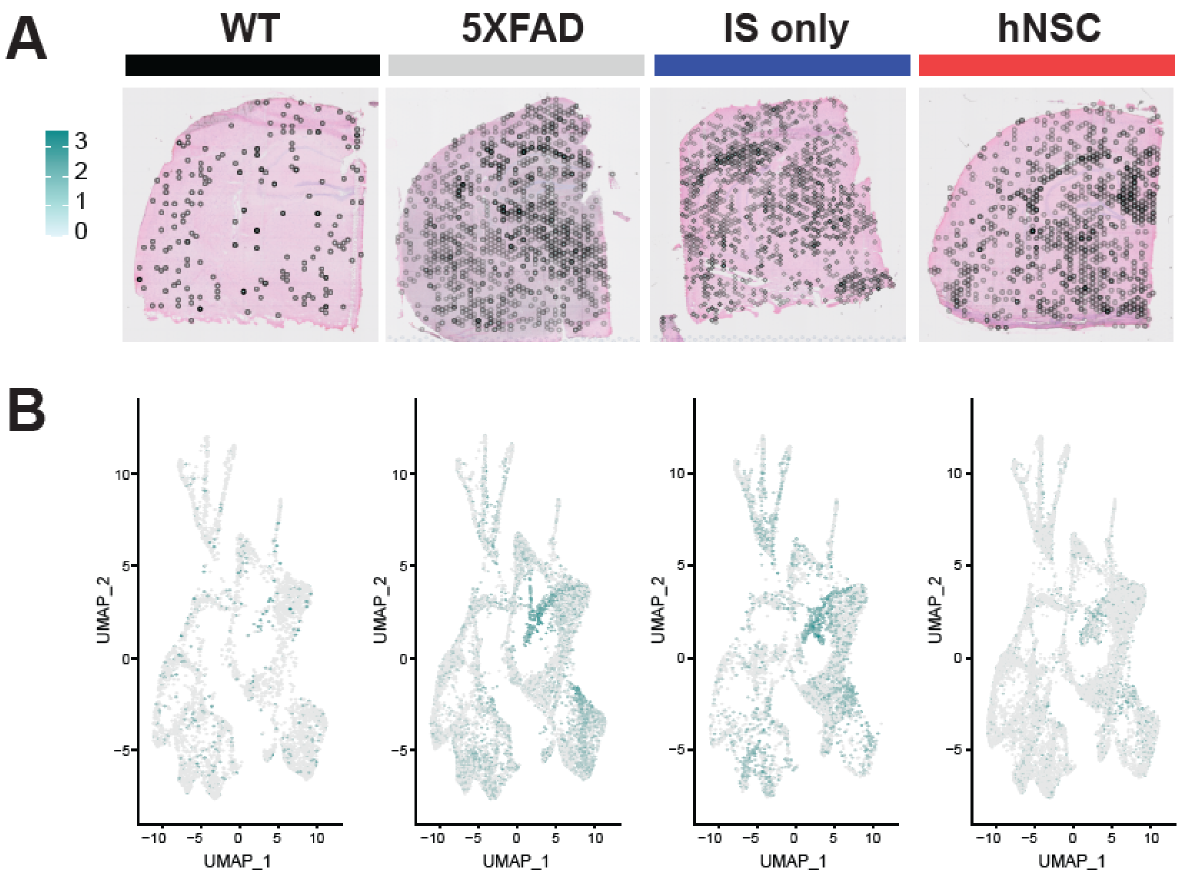

**Figure S9. hNSC transplantation normalizes differentially expressed genes (DEGs) in the hippocampus of 5XFAD mice for 34 weeks.** (**A-B**) Volcano plot of DEGs in (**A**) 5XFAD versus wild-type and (**B**) hNSC versus 5XFAD hippocampus. Upregulated DEGs in green, downregulated DEGs in purple, non-significant or unaffected genes in black. Vertical lines represent average $\log_2$FC= ± 0.6 (FC, fold-change), horizontal line represents $-\log_{10}$(Padj)= $-\log_{10}$(0.05) (Padj, adjusted p-value).

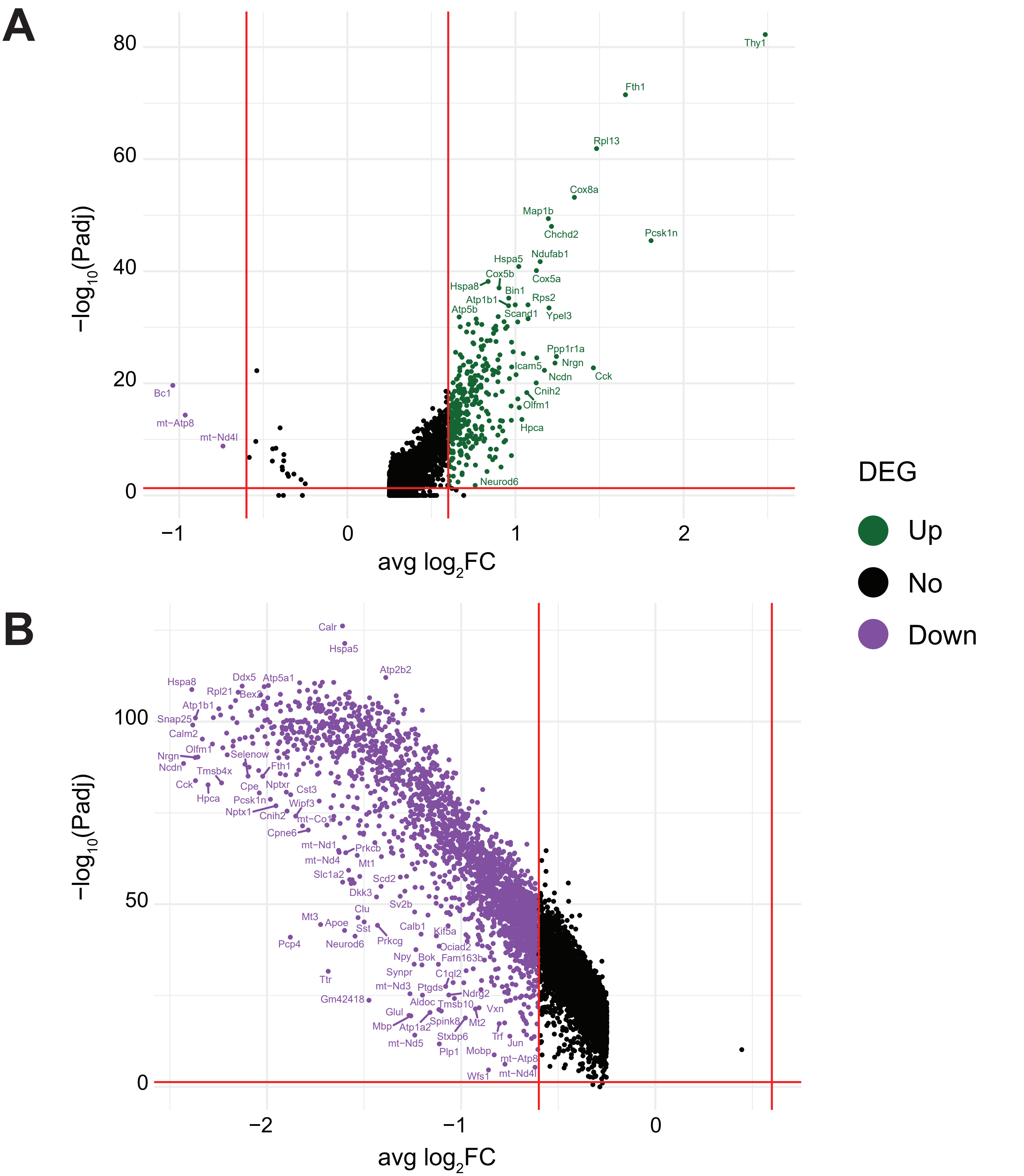

**Figure S10. PIGs expression is highest in microglia.** PIGs feature plot for (**A**) wild-type, (**B**) untreated 5XFAD, (**C**) immunosuppression only (IS only) treated 5XFAD, and (**D**) hNSC injected 5XFAD groups. Plot represents PIGs (x-axis) for various brain areas or cell types (y-axis). PIGs average expression level represented by color scale from less (grey) to higher (purple) expression. PIGs expression is highest in microglia in all groups. Percent expressed represented by dot size from smallest (expressed by none) to largest (100%, expressed by all cells from the cluster). AN, amygdalar nucleus; BNST, bed nucleus of the stria terminalis; CA, ML, cornu ammonis, molecular layer; CA1, PL, cornu ammonis 1, pyramidal layer; CA3, PL, cornu ammonis 1, pyramidal layer; CC, corpus callosum; CP, choroid plexus; DG, GL, dentate gyrus, granule cell layer; DG, ML, dentate gyrus, molecular layer; IC, internal capsule; L1, cortical layer 1; L2/3, cortical layer 2/3; L4, cortical layer 4; L5, cortical layer 5; L6a, L6b, cortical layers 6a and 6b; ZI, zona incerta.

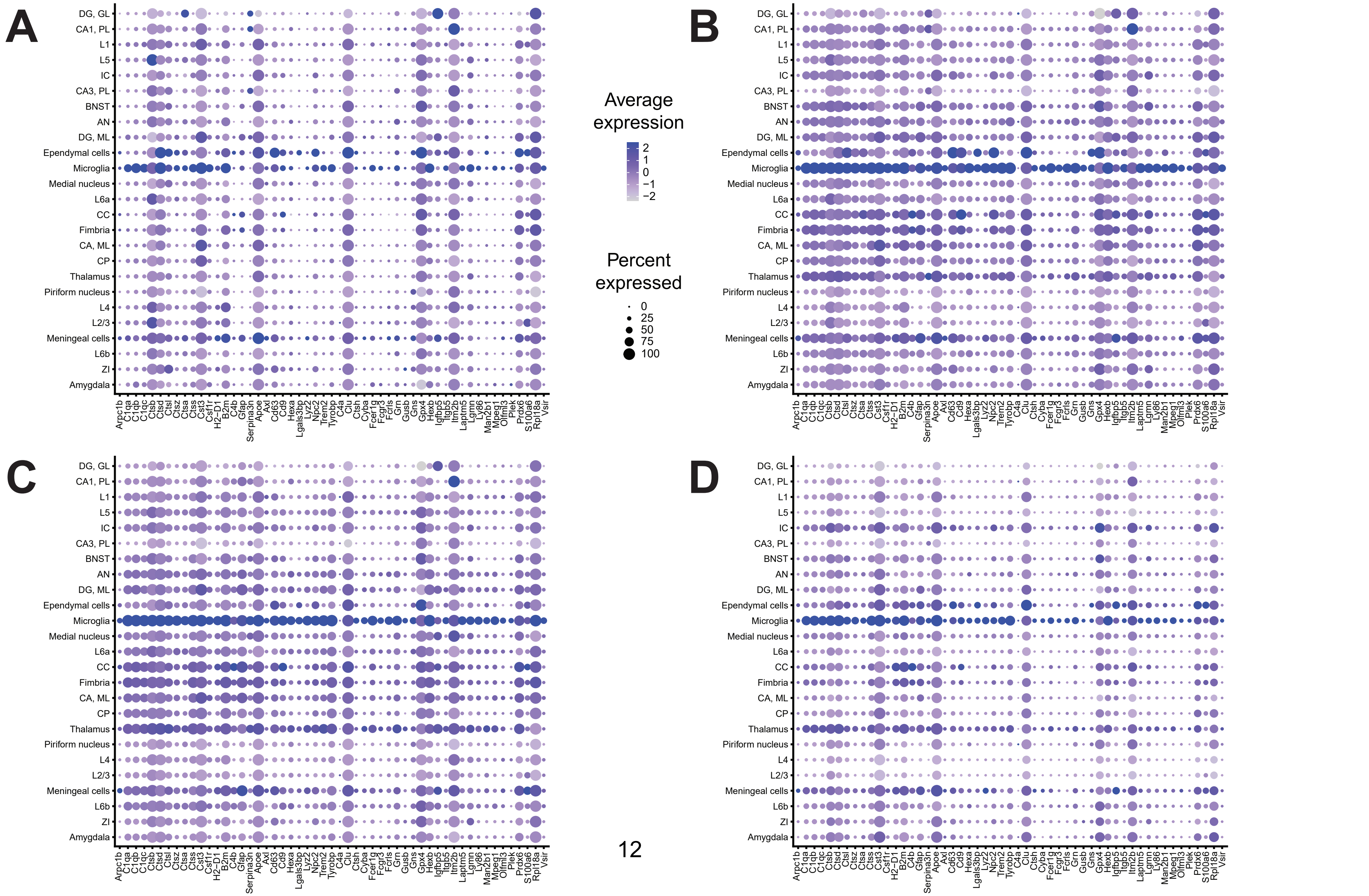

**Figure S11. hNSC transplantation normalizes microglial phenotypes in the brain of 5XFAD mice after 34 weeks.** (**A**) Represenative brain sections of wild-type (WT, black), untreated 5XFAD (grey), immunosuppression only (IS only) treated 5XFAD (IS only, blue), and hNSC injected 5XFAD (red) mice. Homeostatic microglia (HMG; cyan), stage 1 disease-associated microglia (DAMs; green), and stage 2 DAMs (purple) represented by dots. (**B**) Biomarker dot plot of HMG (cyan), stage 1 DAMs (green), and stage 2 DAMs (purple). Plot represents gene biomarkers (x-axis) for various groups (y-axis). Gene average expression level represented by color scale from less (grey) to higher (purple) expression. Percent expressed represented by dot size from smallest (expressed by none) to largest (100%, expressed by all cells from the microglial subtype).

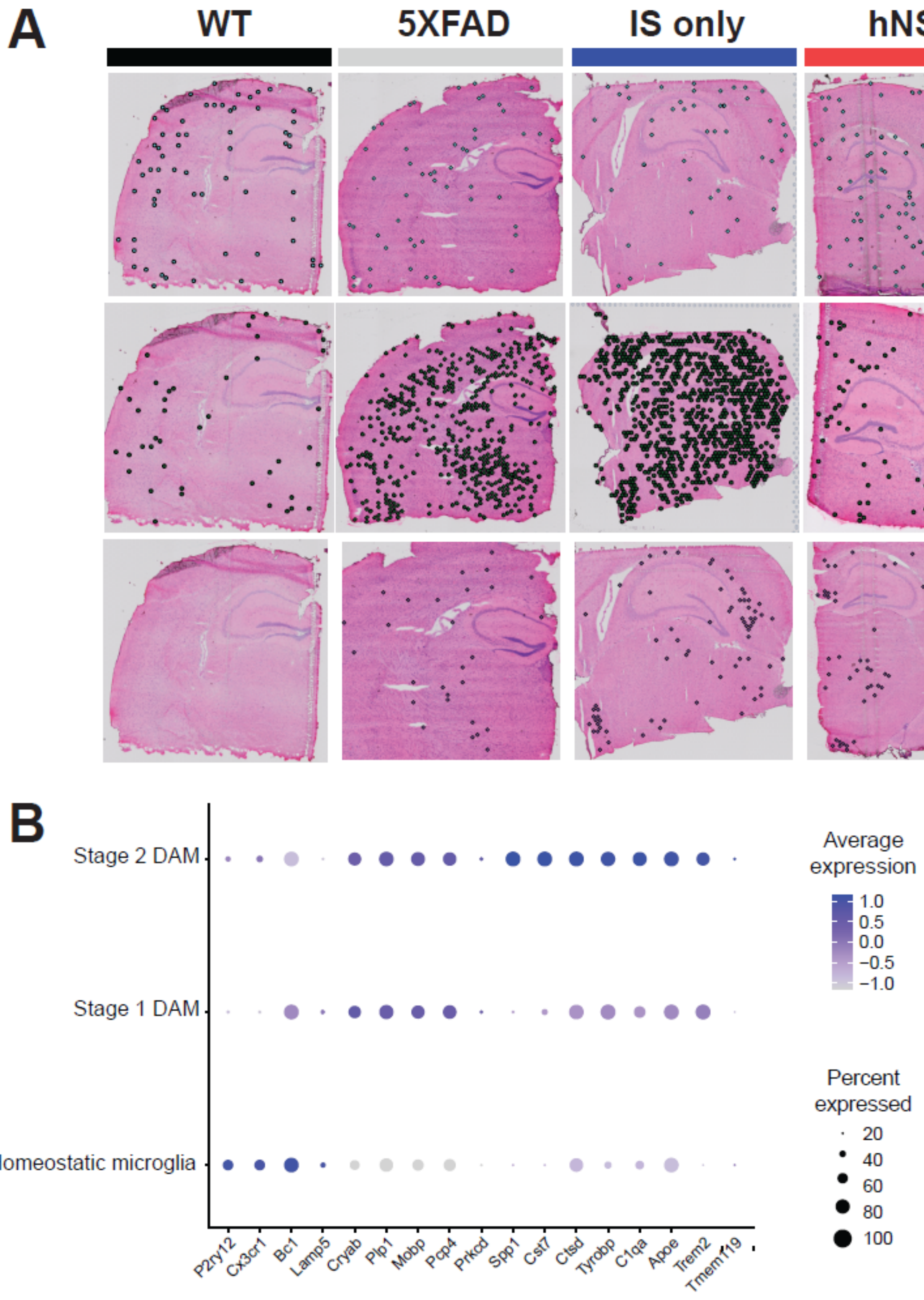

**Figure S12. Intercellular signaling across cell types from WT, 5XFAD, and hNSC brain.** Information flow charts of intercellular signaling across (**A**) wild-type (WT; black) versus untreated 5XFAD (grey) and (**B**) untreated 5XFAD (grey) versus hNSC injected 5XFAD (red) brain cell types generated by CellChat. Vertical dashed lines represent information flow equal in both WT versus 5XFAD and 5XFAD versus hNSC brain cell types. Y-axis labels are colored in the same color as the condition they are significantly upregulated; non-significantly different pathways are labelled in cyan text.

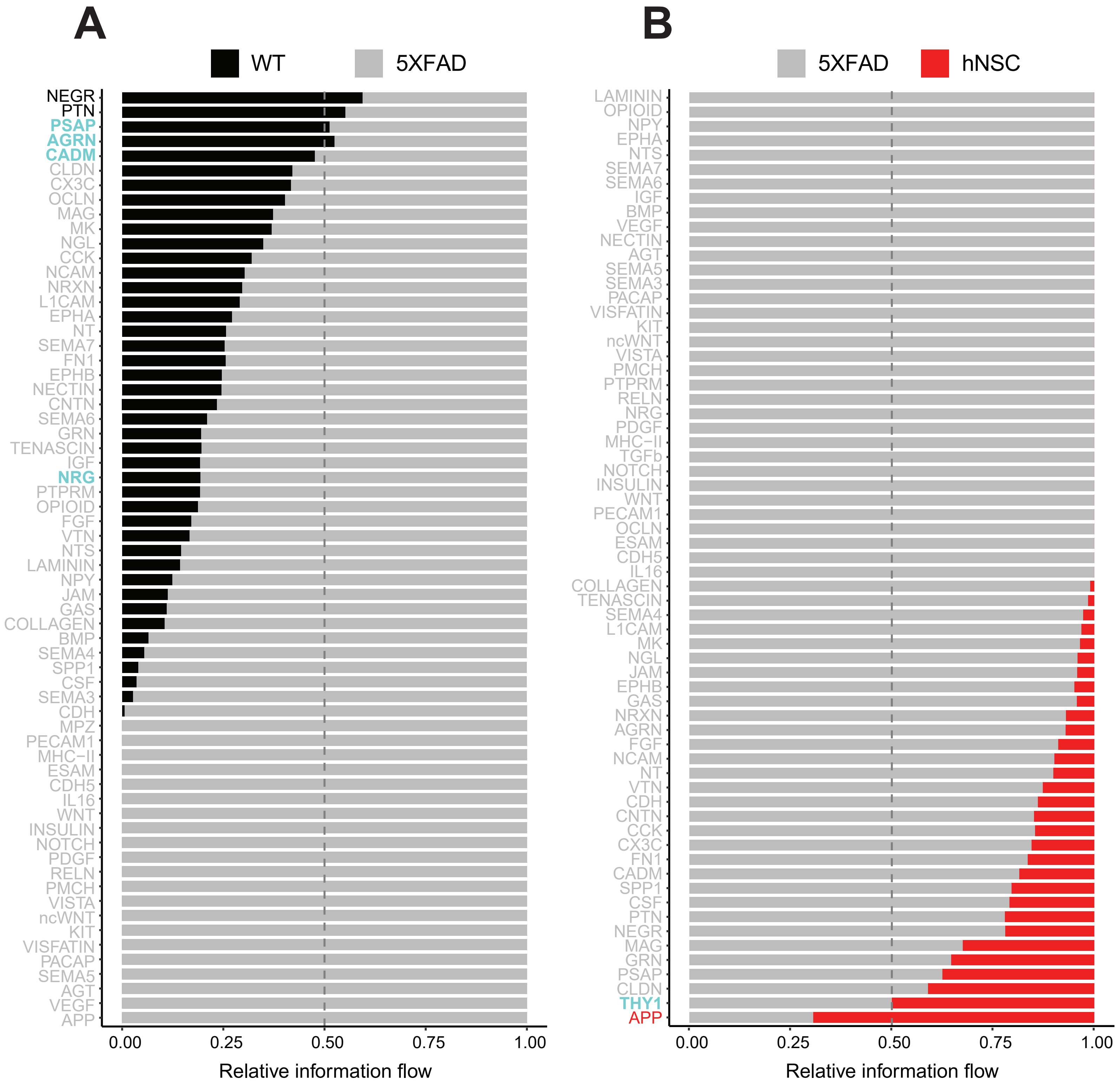

**Figure S13. Pathway enrichment analysis of intercellular signaling from information flow .** Pathway enrichment analysis of intercellular signaling from information flow by (**A**) Kyoto Encyclopedia of Genes and Genomes (KEGG) for 5XFAD versus WT, (**B**) Gene Ontology (GO) for 5XFAD versus WT, (**C**) KEGG for hNSC versus 5XFAD, (**D**) GO for hNSC versus 5XFAD.

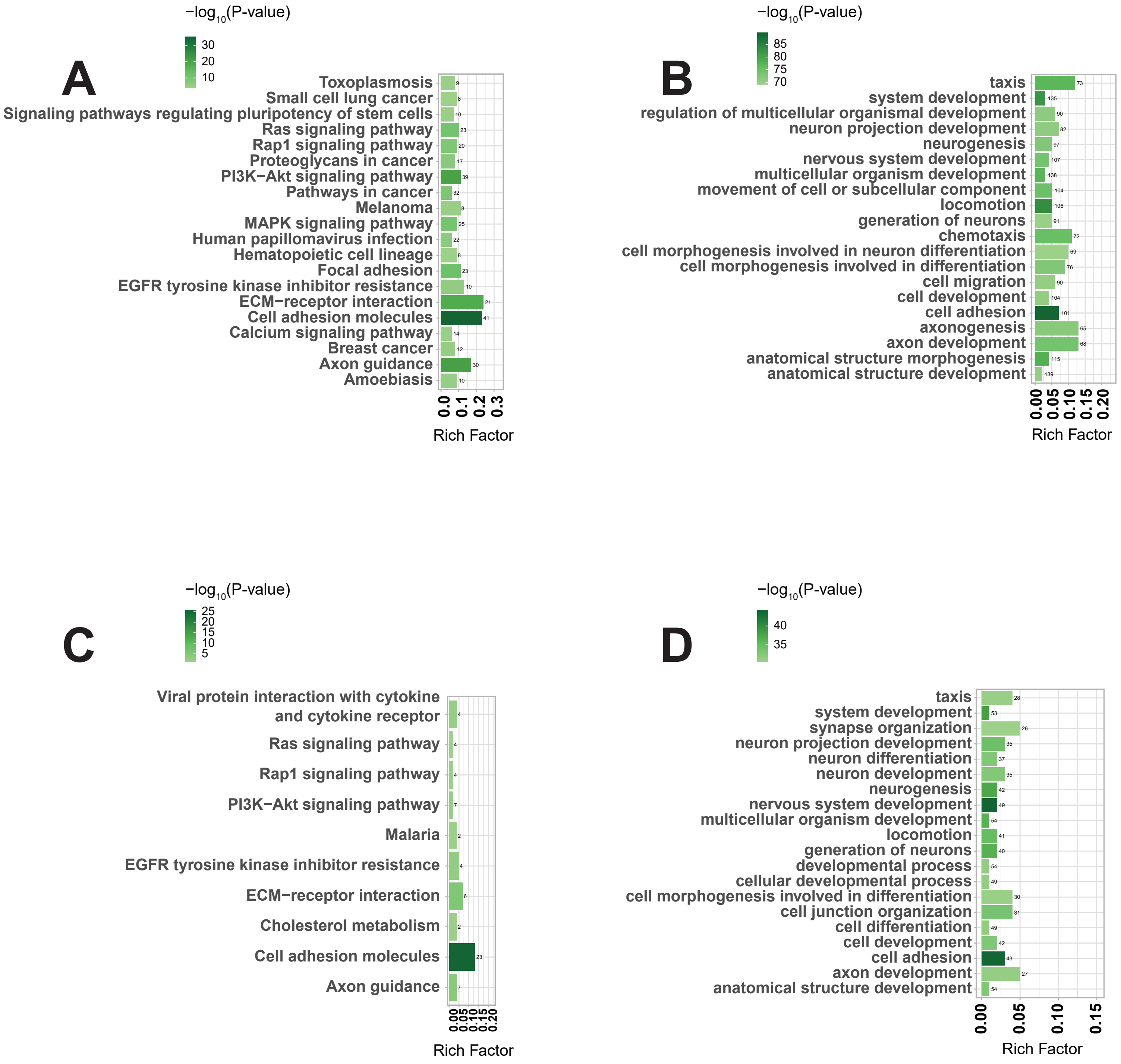

**Figure S14. Ligand-receptor pair interactions between cell types from WT, 5XFAD, and hNSC brains.** Circle plots of ligand-receptor pair interactions between cell types from (**A**) wild-type (WT), (**B**) untreated 5XFAD, (**C**) vehicle injected 5XFAD, and (**D**) hNSC injected 5XFAD brains. Dots represent distinct cell populations; strokes represent communication between distinct cell populations; loops represent signaling within cell populations. Stroke and loop colors reflect the cluster sending the signal, thickness reflects strength of the signaling pair.

BNST, bed nucleus of the stria terminalis; CC, corpus callosum; CPu, caudate putamen; EC, ependymal cells; MC, meningeal cells; MN, medial nucleus; PN, piriform nucleus; ZI, zona incerta.

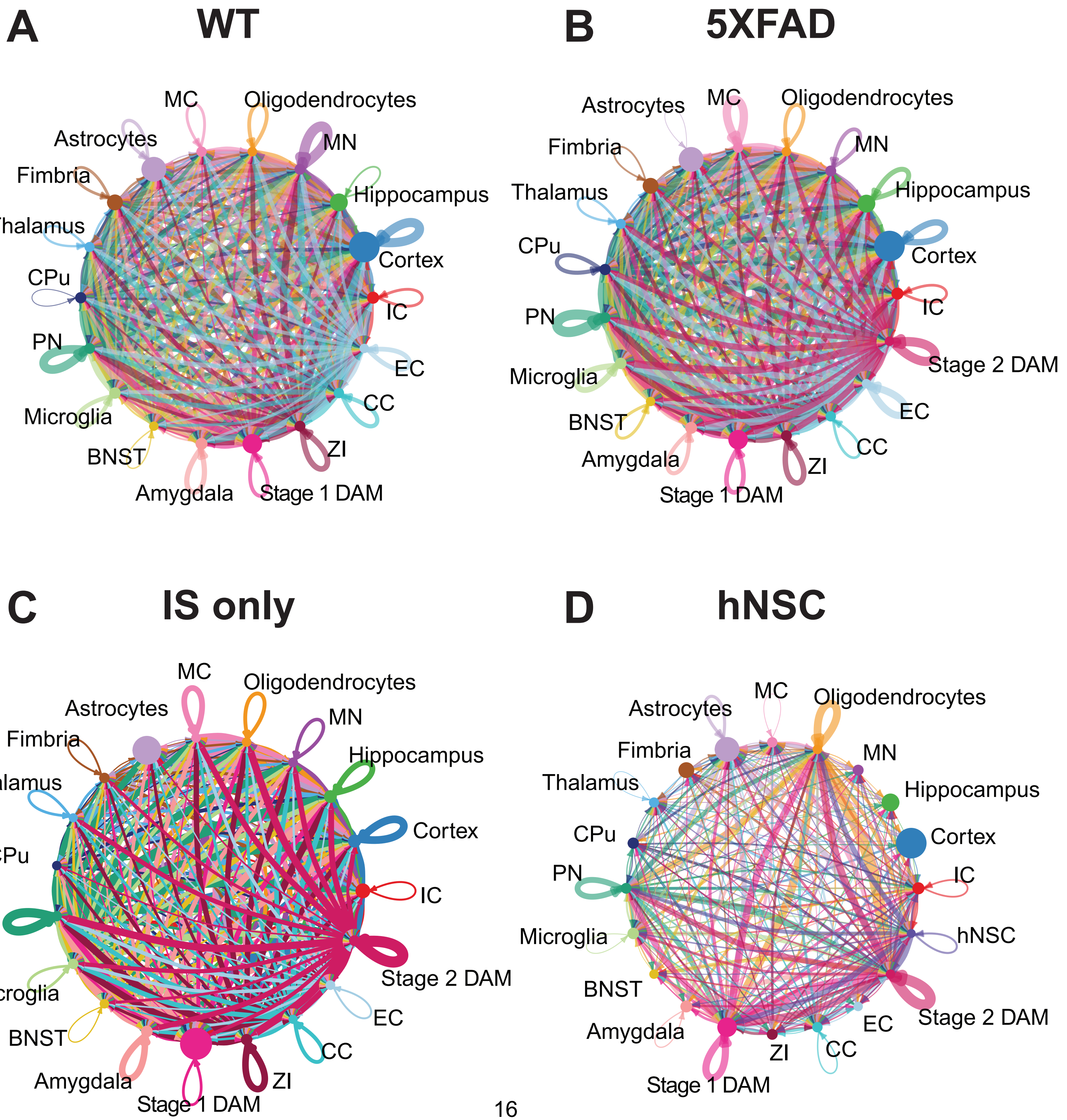

**Figure S15. Pathway enrichment analysis of intercellular signaling between hNSCs and stage 1 and stage 2 DAMs.** Pathway enrichment analysis of intercellular signaling between hNSCs and disease associated microglia (DAMs) by (**A**) Kyoto Encyclopedia of Genes and Genomes (KEGG) for hNSCs versus stage 1 DAMs, (**B**) Gene Ontology (GO) for hNSCs versus stage 1 DAMs, (**C**) KEGG for hNSCs versus stage 2 DAMs, (**D**) GO for hNSCs versus stage 2 DAMs.

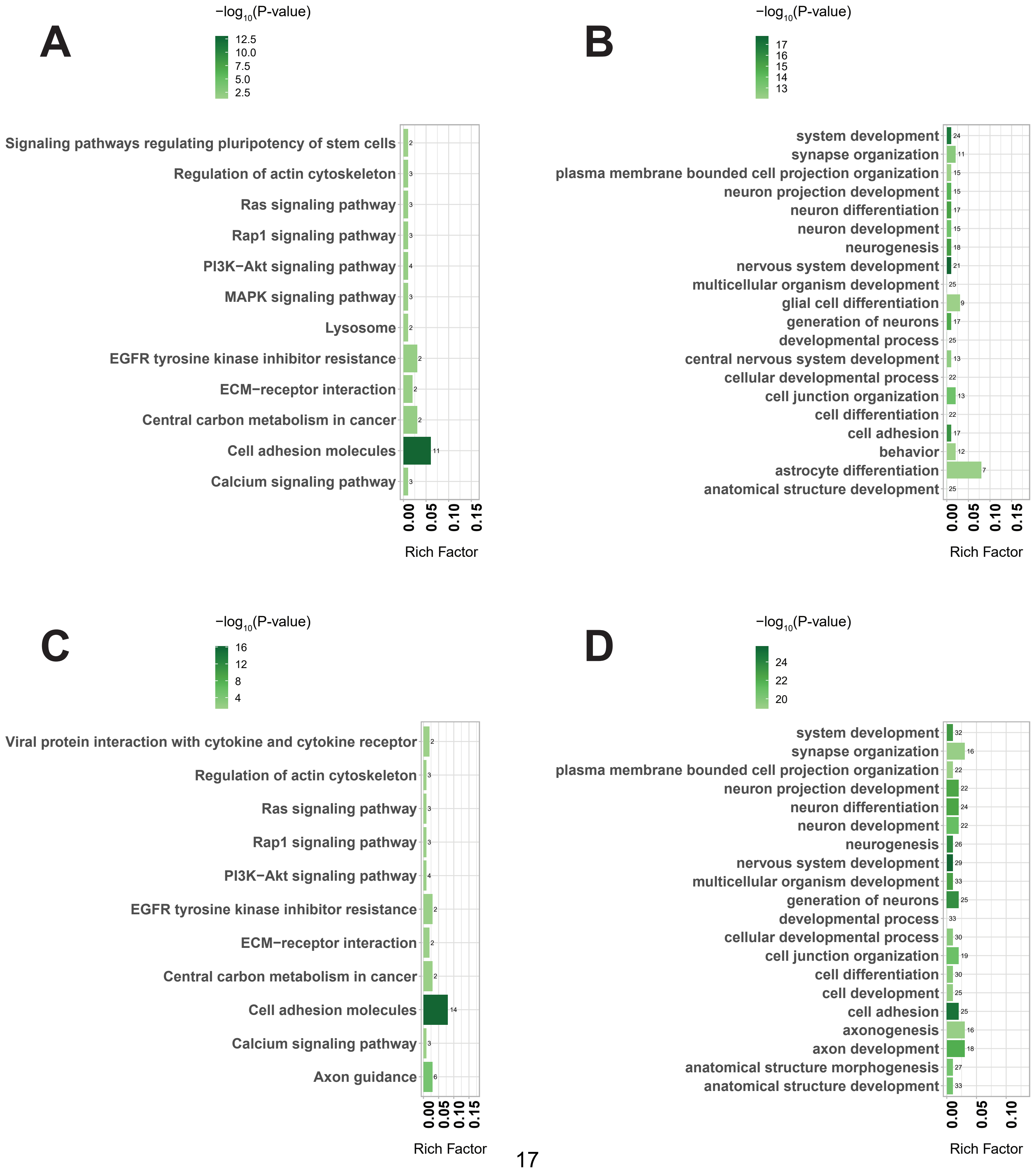

**Figure S16. Differential ligand-receptor pair interactions across cells in 5XFAD versus WT and hNSC versus 5XFAD conditions.** Dot plots of differential ligand-receptor pair interactions between cell types from 5XFAD versus WT that are (**A**) downregulated and (**B**) upregulated. Dot plots of differential ligand-receptor pair interactions between cell types from hNSC versus 5XFAD that are (**C**) downregulated and (**D**) upregulated. Dots represent ligand-receptor signaling (y-axis) between defined cell populations (x-axis; red font, more interactions; blue font, fewer interactions); dot size represents significance value (p-value); dot color represents communication probably from least (blue) to most (red) probable. DAM, stage 1 disease-associated microglia; hippo, hippocampus; MG, microglia; S2DAM, stage 2 disease-associated microglia.

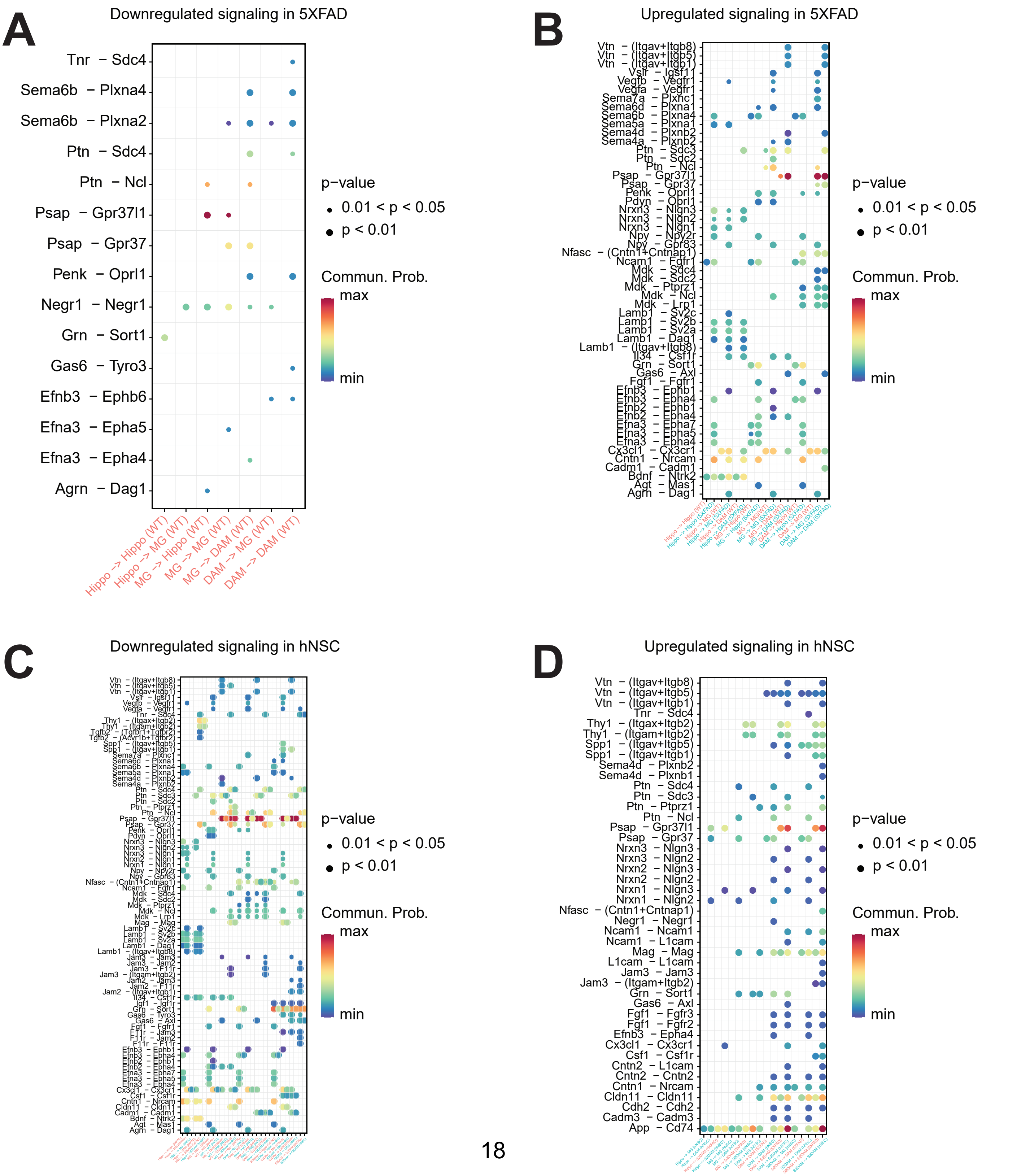

### SUPPLEMENTARY TABLES

#### Table S1. Differentially expressed genes (DEGs) in the whole brain

DEGs in 5XFAD versus wild-type (WT) (grey), immunosuppression only (IS only) treated 5XFAD (IS only, blue), and hNSC versus 5XFAD (red) in the whole brain. DEGs are annotated by gene name, nominal p-value, and Bonferroni-adjusted p-value (Padj). Upregulated DEGs have a positive $\log_2$FC value; downregulated DEGs have a negative $\log_2$FC value. FC, fold-change.

#### Table S2. Differentially expressed genes (DEGs) by cell type

DEGs in 5XFAD versus wild-type (WT), immunosuppression only (IS only) treated 5XFAD versus 5XFAD, and hNSC versus 5XFAD by cell type. DEGs are annotated by gene name, nominal p-value, and Bonferroni-adjusted p-value (Padj). Upregulated DEGs have a positive $\log_2$FC value; downregulated DEGs have a negative $\log_2$FC value. FC, fold-change.

BNST, bed nucleus of the stria terminalis; CA, ML, cornu ammonis, molecular layer; CA1, PL, cornu ammonis 1, pyramidal layer; CA3, PL, cornu ammonis 1, pyramidal layer; CC, corpus callosum; CP, choroid plexus; DG, GL, dentate gyrus, granule cell layer; DG, ML, dentate gyrus, molecular layer; IC, internal capsule; L1, cortical layer 1; L2/3, cortical layer 2/3; L4, cortical layer 4; L5, cortical layer 5; L6a, L6b, cortical layers 6a and 6b; ZI, zona incerta.

#### Table S3. KEGG pathway analysis of DEGs uniquely downregulated by hNSCs in whole brain

Kyoto Encyclopedia of Genes and Genomes (KEGG) pathway analysis of 109 uniquely DEGs downregulated by hNSCs in whole brain. KEGG pathway ID; Term, KEGG pathway name; Annotated, number of DEGs within KEGG pathway; Significant, number of significant DEGs within KEGG pathway; P-value, significance level KEGG pathway; Padj, adjusted significance level KEGG pathway; GeneID, gene name for KEGG pathway DEGs.

#### Table S4. GO pathway analysis of DEGs uniquely downregulated by hNSCs in whole brain

Gene Ontology (GO) pathway analysis of 109 DEGs uniquely downregulated by hNSCs in whole brain. GO pathway ID; Term, GO pathway name; Annotated, number of DEGs within GO pathway; Significant, number of significant DEGs within GO pathway; P-value, significance level GO pathway; Padj, adjusted significance level GO pathway; GeneID, gene name for GO pathway DEGs.

#### Table S5. KEGG pathway analysis of DEGs uniquely downregulated by hNSCs in hippocampus

Kyoto Encyclopedia of Genes and Genomes (KEGG) pathway analysis of 1061 uniquely DEGs downregulated by hNSCs in hippocampus. KEGG pathway ID; Term, KEGG pathway name; Annotated, number of DEGs within KEGG pathway; Significant, number of significant DEGs within KEGG pathway; P-value, significance level KEGG pathway; Padj, adjusted significance level KEGG pathway; GeneID, gene name for KEGG pathway DEGs.

**Table S6. GO pathway analysis of DEGs uniquely downregulated by hNSCs in hippocampus**

Gene Ontology (GO) pathway analysis of 1061 DEGs uniquely downregulated by hNSCs in hippocampus. GO pathway ID; Term, GO pathway name; Annotated, number of DEGs within GO pathway; Significant, number of significant DEGs within GO pathway; P-value, significance level GO pathway; Padj, adjusted significance level GO pathway; GeneID, gene name for GO pathway DEGs.

**Table S7. KEGG and GO pathway analysis of intercellular signaling from information flow in 5XFAD versus WT**

**Sheet 1**: Kyoto Encyclopedia of Genes and Genomes (KEGG) pathway analysis; KEGG pathway ID; Term, KEGG pathway name; Annotated, number of DEGs within KEGG pathway; Significant, number of significant DEGs within KEGG pathway; P-value, significance level KEGG pathway; Padj, adjusted significance level KEGG pathway; GeneID, gene name for KEGG pathway DEGs. **Sheet 2**: Gene Ontology (GO) pathway analysis; GO pathway ID; Term, GO pathway name; Annotated, number of DEGs within GO pathway; Significant, number of significant DEGs within GO pathway; P-value, significance level GO pathway; Padj, adjusted significance level GO pathway; GeneID, gene name for GO pathway DEGs.

**Table S8. KEGG and GO pathway analysis of intercellular signaling from information flow in hNSC versus 5XFAD**

**Sheet 1**: Kyoto Encyclopedia of Genes and Genomes (KEGG) pathway analysis; KEGG pathway ID; Term, KEGG pathway name; Annotated, number of DEGs within KEGG pathway; Significant, number of significant DEGs within KEGG pathway; P-value, significance level KEGG pathway; Padj, adjusted significance level KEGG pathway; GeneID, gene name for KEGG pathway DEGs. **Sheet 2**: Gene Ontology (GO) pathway analysis; GO pathway ID; Term, GO pathway name; Annotated, number of DEGs within GO pathway; Significant, number of significant DEGs within GO pathway; P-value, significance level GO pathway; Padj, adjusted significance level GO pathway; GeneID, gene name for GO pathway DEGs.

**Table S9. Ligand-receptor pairs across all cells in 5XFAD versus WT**

BNST, bed nucleus of the stria terminalis; CA, ML, cornu ammonis, molecular layer; CA1, PL, cornu ammonis 1, pyramidal layer; CA3, PL, cornu ammonis 1, pyramidal layer; CC, corpus callosum; CP, choroid plexus; DG, GL, dentate gyrus, granule cell layer; DG, ML, dentate gyrus, molecular layer; IC, internal capsule; L1, cortical layer 1; L2/3, cortical layer 2/3; L4, cortical layer 4; L5, cortical layer 5; L6a, L6b, cortical layers 6a and 6b; ZI, zona incerta.

**Table S10. Ligand-receptor pairs across all cells in hNSC versus 5XFAD**

BNST, bed nucleus of the stria terminalis; CA, ML, cornu ammonis, molecular layer; CA1, PL, cornu ammonis 1, pyramidal layer; CA3, PL, cornu ammonis 1, pyramidal layer; CC, corpus callosum; CP, choroid plexus; DG, GL, dentate gyrus, granule cell layer; DG, ML, dentate gyrus, molecular layer; IC, internal capsule; L1, cortical layer 1; L2/3, cortical layer 2/3; L4, cortical layer 4; L5, cortical layer 5; L6a, L6b, cortical layers 6a and 6b; ZI, zona incerta.

**Table S11. Ligand-receptor pairs in hNSCs versus stage 1 DAMs and stage 2 DAMs**

**Table S12. KEGG and GO pathway analysis of intercellular signaling between hNSC and stage 1 and stage 2 DAMs**

**Sheet 1**: Kyoto Encyclopedia of Genes and Genomes (KEGG) pathway analysis for intercellular signaling from hNSCs to stage 1 DAMs; **Sheet 2**: KEGG for intercellular signaling from hNSCs to stage 2 DAMs.

KEGG pathway ID; Term, KEGG pathway name; Annotated, number of DEGs within KEGG pathway; Significant, number of significant DEGs within KEGG pathway; P-value, significance level KEGG pathway; Padj, adjusted significance level KEGG pathway; GeneID, gene name for KEGG pathway DEGs.

**Sheet 3**: Gene Ontology (GO) pathway analysis for intercellular signaling from hNSCs to stage 1 DAMs; **Sheet 4**: GO for intercellular signaling from hNSCs to stage 2 DAMs.

GO pathway ID; Term, GO pathway name; Annotated, number of DEGs within GO pathway; Significant, number of significant DEGs within GO pathway; P-value, significance level GO pathway; Padj, adjusted significance level GO pathway; GeneID, gene name for GO pathway DEGs.

**Table S13. Ligand-receptor pairs stage 1 DAMs and stage 2 DAMs versus hNSCs**

**Table S14. Differentially expressed genes (DEGs) in the whole brain by linear mixed effects model**

**Table S15. Differentially expressed genes (DEGs) in the hippocampus by linear mixed effects model**
